# Genome sequencing analysis identifies new loci associated with Lewy body dementia and provides insights into the complex genetic architecture

**DOI:** 10.1101/2020.07.06.185066

**Authors:** Ruth Chia, Marya S. Sabir, Sara Bandres-Ciga, Sara Saez-Atienzar, Regina H. Reynolds, Emil Gustavsson, Ronald L. Walton, Sarah Ahmed, Coralie Viollet, Jinhui Ding, Mary B. Makarious, Monica Diez-Fairen, Makayla K. Portley, Zalak Shah, Yevgeniya Abramzon, Dena G. Hernandez, Cornelis Blauwendraat, David J. Stone, John Eicher, Laura Parkkinen, Olaf Ansorge, Lorraine Clark, Lawrence S. Honig, Karen Marder, Afina Lemstra, Peter St George-Hyslop, Elisabet Londos, Kevin Morgan, Tammaryn Lashley, Thomas T. Warner, Zane Jaunmuktane, Douglas Galasko, Isabel Santana, Pentti J. Tienari, Liisa Myllykangas, Minna Oinas, Nigel J. Cairns, John C. Morris, Glenda M. Halliday, Vivianna M. Van Deerlin, John Q. Trojanowski, Maurizio Grassano, Andrea Calvo, Gabriele Mora, Antonio Canosa, Gianluca Floris, Ryan C. Bohannan, Francesca Brett, Ziv Gan-Or, Joshua T. Geiger, Anni Moore, Patrick May, Rejko Krüger, David Goldstein, Grisel Lopez, Nahid Tayebi, Ellen Sidransky, the Fox Investigation for New Discovery of Biomarkers; The American Genome Center; Lucy Norcliffe-Kaufmann, Jose-Alberto Palma, Horacio Kaufmann, Vikram Shakkottai, Matthew Perkins, Kathy L. Newell, Thomas Gasser, Claudia Schulte, Francesco Landi, Erika Salvi, Daniele Cusi, Eliezer Masliah, Ronald C. Kim, Chad A. Caraway, Ed Monuki, Maura Brunetti, Ted M. Dawson, Liana S. Rosenthal, Marilyn S. Albert, Olga Pletnikova, Juan C. Troncoso, Margaret E. Flanagan, Qinwen Mao, Eileen H. Bigio, Eloy Rodríguez-Rodríguez, Jon Infante, Carmen Lage, Isabel González-Aramburu, Pascual Sanchez-Juan, Bernardino Ghetti, Julia Keith, Sandra E. Black, Mario Masellis, Ekaterina Rogaeva, Charles Duyckaerts, Alexis Brice, Suzanne Lesage, Georgia Xiromerisiou, Matthew J. Barrett, Bension S. Tilley, Steve Gentleman, Giancarlo Logroscino, Geidy E. Serrano, Thomas G. Beach, Ian G. McKeith, Alan J. Thomas, Johannes Attems, Christopher M. Morris, Laura Palmer, Seth Love, Claire Troakes, Safa Al-Sarraj, Angela K. Hodges, Dag Aarsland, Gregory Klein, Scott M. Kaiser, Randy Woltjer, Pau Pastor, Lynn M. Bekris, James Leverenz, Lilah M. Besser, Amanda Kuzma, Alan E. Renton, Alison Goate, David A. Bennett, Clemens R. Scherzer, Huw R. Morris, Raffaele Ferrari, Diego Albani, Stuart Pickering- Brown, Kelley Faber, Walter Kukull, Estrella Morenas-Rodriguez, Alberto Lleó, Juan Fortea, Daniel Alcolea, Jordi Clarimon, Michael A. Nalls, Luigi Ferrucci, Susan M. Resnick, Toshiko Tanaka, Tatiana M. Foroud, Neill R. Graff-Radford, Zbigniew K. Wszolek, Tanis Ferman, Bradley F. Boeve, John A. Hardy, Eric Topol, Ali Torkamani, Andrew B. Singleton, Mina Ryten, Dennis Dickson, Adriano Chiò, Owen A. Ross, J. Raphael Gibbs, Clifton L. Dalgard, Bryan J. Traynor, Sonja W. Scholz

## Abstract

The genetic basis of Lewy body dementia (LBD) is not well understood. Here, we performed whole-genome sequencing in large cohorts of LBD cases and neurologically healthy controls to study the genetic architecture of this understudied form of dementia and to generate a resource for the scientific community. Genome-wide association analysis identified five independent risk loci, whereas genome-wide gene-aggregation tests implicated mutations in the gene *GBA*. Genetic risk scores demonstrate that LBD shares risk profiles and pathways with Alzheimer’s and Parkinson’s disease, providing a deeper molecular understanding of the complex genetic architecture of this age-related neurodegenerative condition.

## Introduction

Lewy body dementia (LBD) is a clinically heterogeneous neurodegenerative disease characterized by progressive cognitive decline, parkinsonism, and visual hallucinations^1^. There are no effective disease-modifying treatments available to slow disease progression, and current therapy is limited to symptomatic and supportive care. At postmortem, the disorder is distinguished by the widespread cortical and limbic deposition of α-synuclein protein in the form of Lewy bodies that are also a hallmark feature of Parkinson’s disease. The vast majority of LBD patients additionally exhibit Alzheimer’s disease co-pathology^2^. These neuropathological observations have led to the, as yet unproven hypothesis that LBD lies on a disease continuum between Parkinson’s disease and Alzheimer’s disease^3^. Though relatively common in the community, with an estimated 1.4 million prevalent cases in the United States^4^, the genetic contributions to this underserved condition are poorly understood.

The rapid advances in genome sequencing technologies offer unprecedented opportunities to identify and characterize disease-associated genetic variation. Here, we performed whole-genome sequencing in a cohort of 2,981 patients diagnosed with LBD and 4,391 neurologically healthy subjects. We analyzed these data using a genome-wide association study (GWAS) approach and gene aggregation tests, and we modeled the relative contributions of Alzheimer’s disease and Parkinson’s disease risk variants to this fatal neurodegenerative disease (see Fig. 1 for an analysis overview). Additionally, we created a resource for the scientific community to mine for new insights into the genetic etiology of LBD and to expedite the development of targeted therapeutics.

**Fig. 1.**
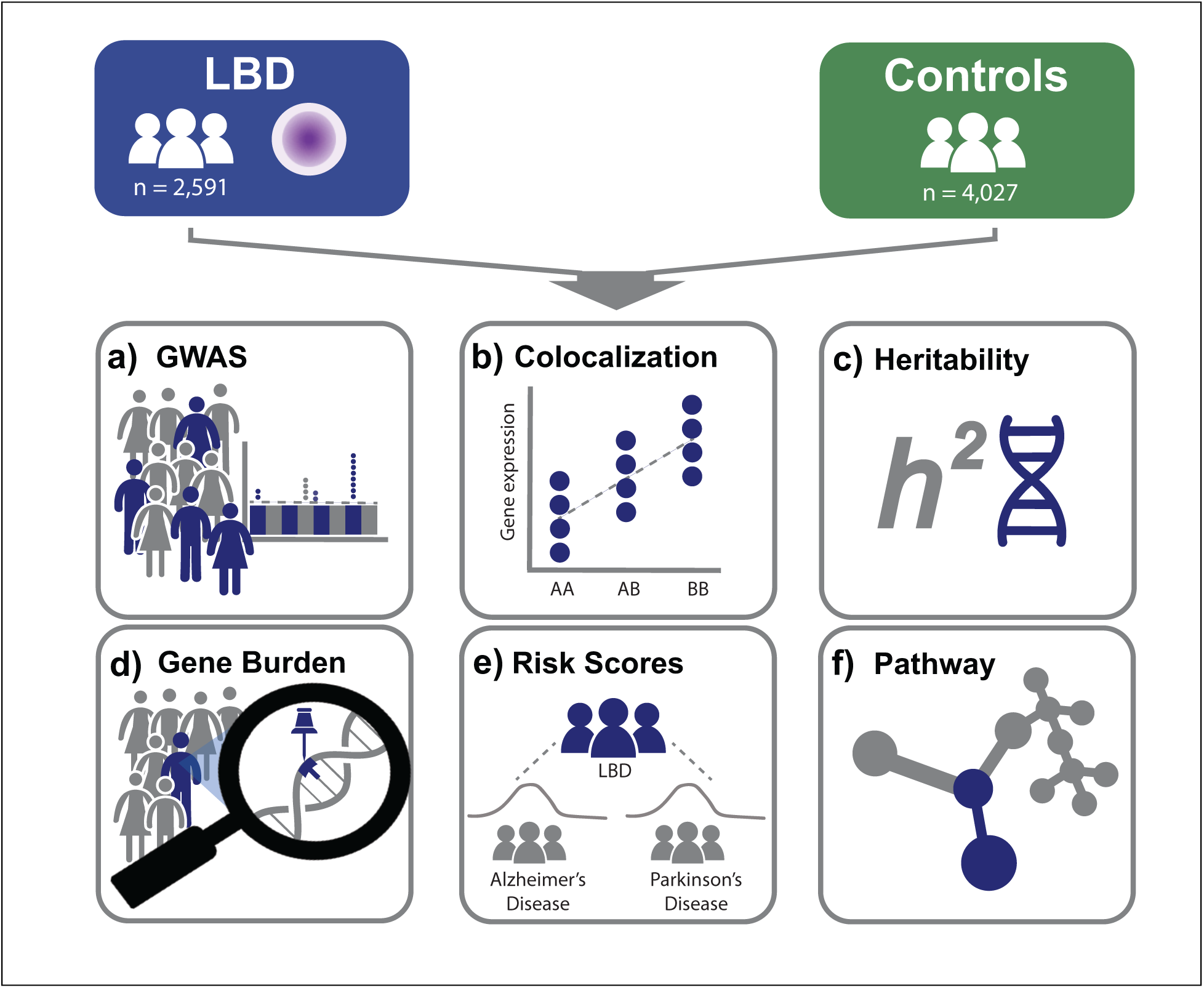
Analysis workflow. Schematic illustration of the analytical workflow.

## Results

### Genome-wide association analysis identifies new loci associated with LBD

After quality control, whole-genome sequence data from 2,591 patients diagnosed with LBD and 4,027 neurologically healthy subjects were available for study. Participants were recruited across 44 institutions/consortia and were diagnosed according to established consensus criteria. Using a GWAS approach, we identified five loci that surpassed the genome-wide significance threshold (Table 1, Fig. 2a). Three of these signals were located at known LBD risk loci within the genes *GBA, APOE,* and *SNCA*^5–8^. The remaining GWAS signals in *BIN1* and *TMEM175* represented novel LBD risk loci. Notably, these loci have been implicated in other age-related neurodegenerative diseases, including Alzheimer’s disease (*BIN1*) and Parkinson’s disease (*TMEM175*)^9, 10^. Conditional analyses detected a second signal at the *APOE* locus (Supplementary Fig. 1 for regional association plots, Supplementary Fig. 2 for conditional association analyses). Subanalysis GWAS of pathologically defined LBD cases only versus control subjects identified the same risk loci (Fig. 2b). Finally, we replicated each of the observed risk loci in an independent sample of 970 European-ancestry LBD cases and 8,928 control subjects (Table 1).

**Fig. 2.**
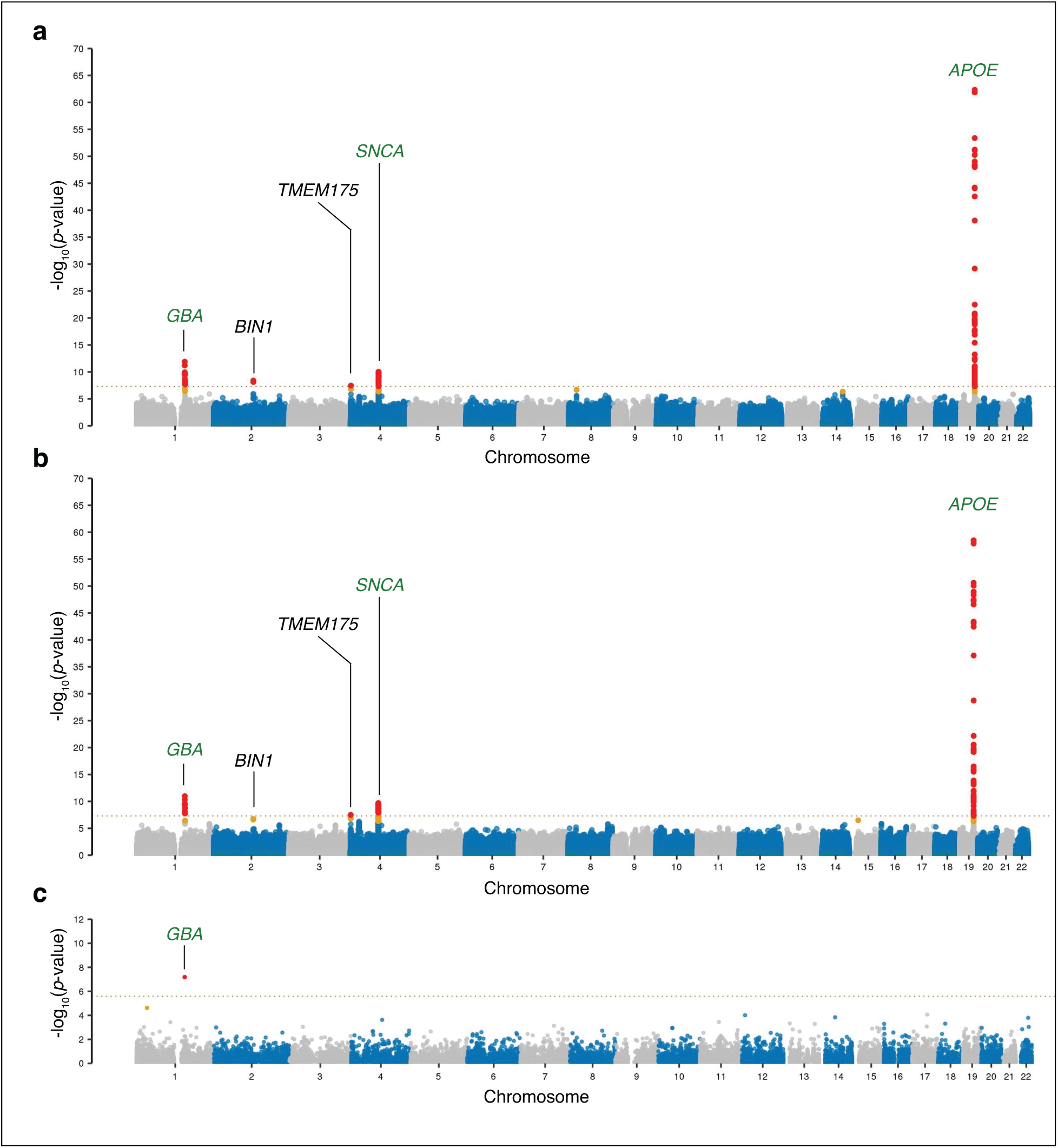
Genome-wide representation of common and rare variant associations in LBD. Manhattan plots depicting **a**, the GWAS results (n = 2,591 cases and 4,027 controls; MAF >1%), **b**, the GWAS subanalysis of pathologically confirmed LBD cases only (n = 1,789) versus controls (n = 4,027), and **c**, gene-based genome-wide SKAT-O test associations of rare missense and loss-of-function variants (MAF ≤ 1%). The x-axis denotes the chromosomal position for all 22 autosomes in hg38, and the y-axis indicates the association *p*-values on a -log_10_ scale. Each dot in **a**, and **b**, indicates a single-nucleotide variant or indel, while each dot in **c**, corresponds to a gene. Red dots highlight genome-wide significant signals, while suggestive variants are indicated with orange dots. A dashed line shows the conservative Bonferroni threshold for genome-wide significance. For **a**, and **b**, the gene with the closest proximity to the top variant at each significant locus is listed. Green font was used to highlight known LBD risk loci, while black font indicates novel association signals.

**Table 1.**
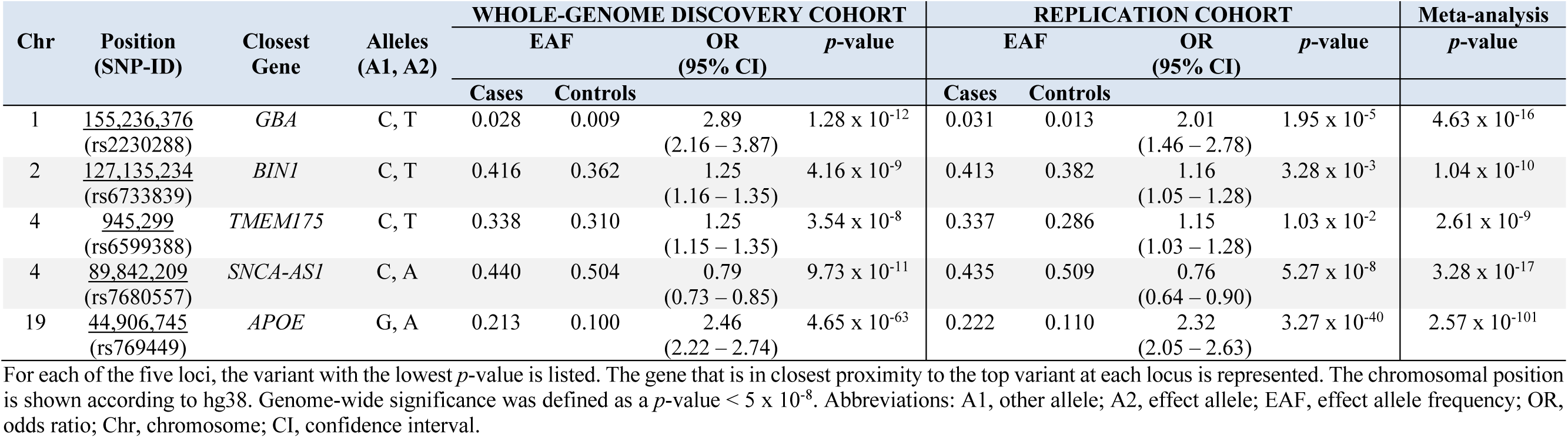
Genome-wide significant association signals in LBD GWAS

### Gene-level aggregation testing identifies *GBA* as a pleomorphic risk gene

The significant loci from our GWAS explained only a small fraction (1%) of the conservatively estimated narrow-sense heritability of LBD of 10.81% (95% confidence interval [CI]: 8.28% – 13.32%, *p*-value = 9.17 × 10^-4^). To explore whether rare variants contribute to the remaining risk of LBD, we performed gene-level sequence kernel association – optimized (SKAT-O) tests of missense and loss-of-function mutations with a minor allele frequency (MAF) threshold ≤ 1% across the genome^11^. This rare variant analysis identified *GBA* as associated with LBD (Fig. 2c). *GBA*, encoding the lysosomal enzyme glucocerebrosidase, is a known pleomorphic risk gene of LBD and Parkinson’s disease^5, 12, 13^, and our rare and common variant analyses confirm a prominent role of this gene in the pathogenesis of Lewy body diseases.

### Functional inferences from colocalization and gene expression analyses

Most GWAS loci are thought to operate through the regulation of gene expression^14, 15^. Thus, we performed a colocalization analysis to determine whether a shared causal variant drives association signals for LBD risk and gene expression. Expression quantitative trait loci (eQTLs) were obtained from eQTLGen and PsychENCODE^16, 17^, the largest available human blood and brain eQTL datasets. We found evidence of colocalization between the *TMEM175* locus and an eQTL regulating *TMEM175* expression in blood (posterior probability for H_4_ (PPH4) = 0.99; Fig. 3a; Supplementary Table 1). There was also colocalization between the association signal at the *SNCA* locus and an eQTL regulating *SNCA-AS1* expression in the brain (PPH4 = 0.96; Fig. 3b; Supplementary Table 1). Interestingly, the index variant at the *SNCA* locus was located within the *SNCA-AS1* gene, which overlaps with the 5’-end of *SNCA* and encodes a long noncoding antisense RNA known to regulate *SNCA* expression. Sensitivity analyses confirmed that these colocalizations were robust to changes in the prior probability of a variant associating with both traits (Supplementary Fig. 3).

**Fig. 3.**
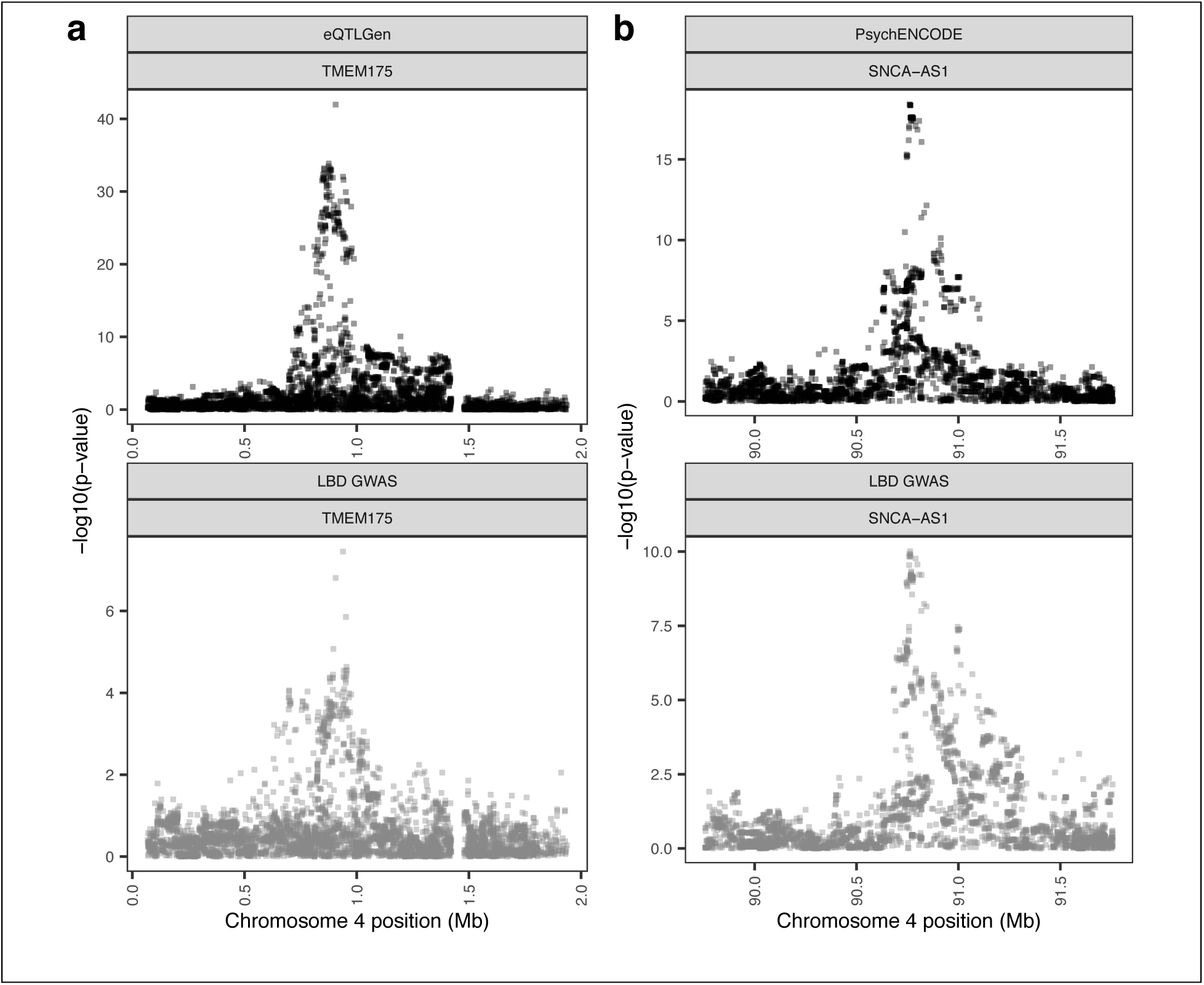
Regional association plots for eQTL and LBD GWAS colocalizations. Regional association plots for eQTLs (upper pane) and LBD GWAS signals (lower pane) in the regions surrounding **a**, *TMEM175* (PPH4 = 0.99) and **b**, *SNCA-AS1* (PPH4 = 0.96). The x-axis denotes the chromosomal position in hg19, and the y-axis indicates the association *p*-values on a -log_10_ scale.

**Fig. 4.**
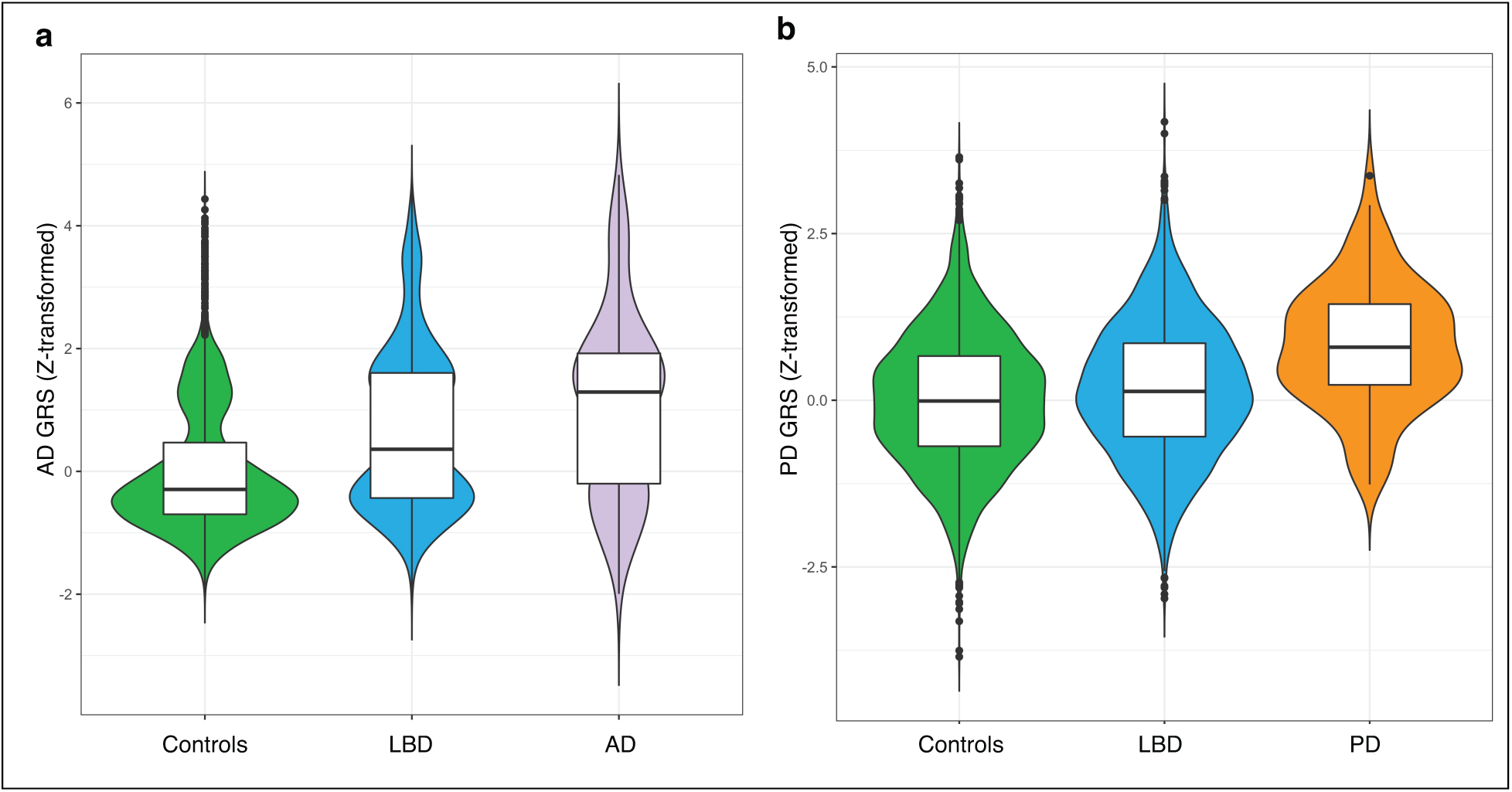
Genetic risk scores from Alzheimer’s disease and Parkinson’s disease GWAS studies illustrate intersecting molecular genetic risk profiles with LBD. Alzheimer’s disease and Parkinson’s disease genetic risk scores predict risk for LBD and highlight overlapping molecular risk profiles. **a**, Violin plots comparing z-transformed Alzheimer’s disease genetic risk score distributions in LBD cases, controls, and 100 random Alzheimer’s disease cases, while **b**, shows the z-transformed Parkinson’s disease genetic risk score distributions for LBD cases, controls, and 100 random Parkinson’s disease cases. The center line of each violin plot is the median, the box limits depict the interquartile range, and whiskers correspond to the 1.5x interquartile range. Abbreviations: GRS, genetic risk score; AD, Alzheimer’s disease; PD, Parkinson’s disease.

We interrogated the effect of each SNP in the region surrounding *SNCA-AS1* on LBD risk using our GWAS data and *SNCA-AS1* expression using the PsychENCODE data (Supplementary Fig. 4a). All genome-wide significant risk SNPs in the locus had a negative beta coefficient, while the shared *SNCA-AS1* eQTL had a positive beta coefficient. This negative correlation suggested that increased *SNCA-AS1* expression is associated with reduced LBD risk (Spearman’s rho = -0.42; *p*-value = 0.0012; Supplementary Fig. 4b).

Analysis of human bulk-tissue RNA-sequencing data from the Genotype-Tissue Expression (GTEx) consortium and single-nucleus RNA-sequencing data of the medial temporal gyrus from the Allen Institute of Brain Science ^18, 19^ demonstrated that *TMEM175* is ubiquitously expressed, whereas *SNCA-AS1* is predominantly expressed in brain tissue (Supplementary Fig. 5a; Supplementary Table 2). At the cellular level, *TMEM175* is highly expressed in oligodendrocyte progenitor cells, while *SNCA-AS1* demonstrates neuronal specificity (Supplementary Fig. 5b; Supplementary Table 2). *SNCA* and *SNCA-AS1* share a similar, though not identical, tissue expression profile (Supplementary Fig. 6).

### LBD risk overlaps with risk profiles of Alzheimer’s disease and Parkinson’s disease

We leveraged our whole-genome sequence data to explore the etiological relationship between Alzheimer’s disease, Parkinson’s disease, and LBD. To do this, we applied genetic risk scores derived from large-scale GWASes in Alzheimer’s and Parkinson’s disease to individual-level genetic data from our LBD case-control cohort^20, 21^. We tested the associations of the Alzheimer’s and Parkinson’s disease genetic risk scores with LBD disease status, and with age at death, age at onset, and the duration of illness observed among the LBD cases.

Patients diagnosed with LBD had a higher genetic risk for developing both Alzheimer’s disease (odds ratio [OR] = 1.66 per standard deviation of Alzheimer’s disease genetic risk, 95% CI = 1.58 - 1.74, *p*-value < 2 × 10^-16^, Fig. 5a) and Parkinson’s disease (OR = 1.20, 95% CI =1.14 - 1.26, *p*-value = 4.34 × 10^-12^, Fig. 5b). These risk scores remained significant after adjusting for genes that substantially contribute to Alzheimer’s disease (model after adjustment for *APOE*: OR = 1.53, 95% CI = 1.37 - 1.72, *p*-value = 3.29 × 10^-14^) and Parkinson’s disease heritable risk (model after adjustment for *GBA, SNCA*, and *LRRK2*: OR = 1.26, 95% CI = 1.19 - 1.34, *p*-value = 5.91 × 10^-14^). The Alzheimer’s disease genetic risk score was also found to be significantly associated with an earlier age of death in LBD (β = -1.77 years per standard deviation increase in the genetic risk score from the population mean, standard error [SE] = 0.19, *p*-value < 2 × 10^-16^) and shorter disease duration (β = -0.90 years, SE = 0.27, *p*-value = 0.0007). In contrast, the Parkinson’s disease genetic risk score was associated with an earlier age at onset among patients diagnosed with LBD (β = -0.98, SE = 0.28, *p*-value = 0.00045), indicating that higher Parkinson’s disease risk is associated with earlier age at onset in LBD. We found no evidence of interaction between the genetic risk scores of Alzheimer’s disease and Parkinson’s disease in the LBD cohort (OR = 0.99, 95 % CI = 0.95 - 1.03, *p*-value = 0.59), implying that Alzheimer’s disease and Parkinson’s disease risk variants are independently associated with LBD risk.

**Fig. 5.**
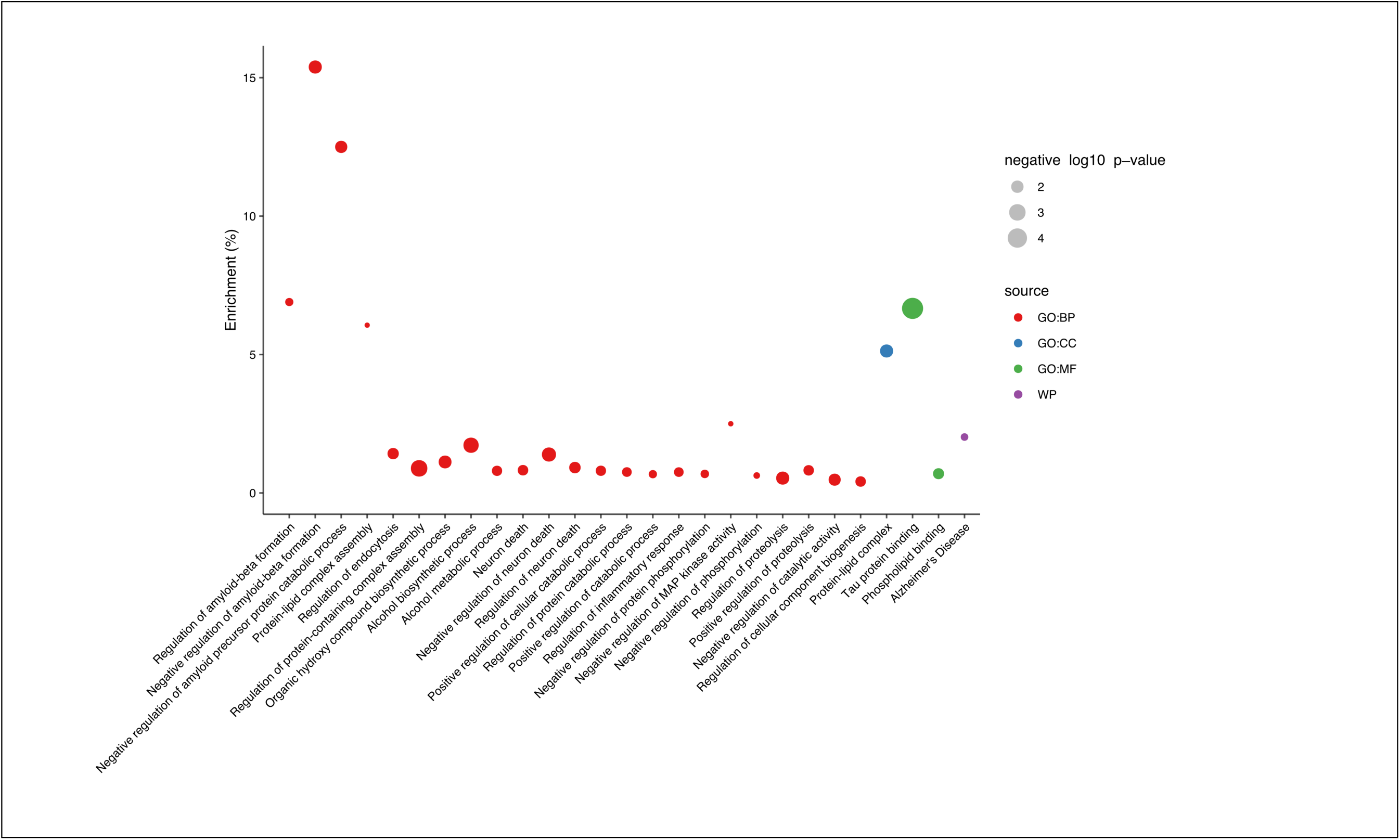
Insights into LBD pathways based on polygenic risk score enrichment analysis. Functional enrichment analyses of the LBD polygenic risk scores. The x-axis corresponds to the enrichment category in LBD cases compared to controls, and the y-axis shows the enrichment percentages of significant associations after multiple testing correction. The enrichment percentage refers to the percentage of input genes/variants that are within in a given pathway. Significant gene ontology (GO) enrichments for biological processes (BP, red), cellular functions (CC, green), molecular processes (MP, blue), and pathways from WikiPathways (WP, purple) are shown. The size of each respective dot indicates the *p*-values on a -log_10_ scale.

### Enrichment analysis identifies pathways involved in LBD

Pathway enrichment analysis of LBD, using a polygenic risk score based on the GWAS risk variants, found several significantly enriched gene ontology processes associated with LBD (Fig. 6). These related to the *regulation of amyloid-beta formation* (adjusted *p*-value = 0.04)*, regulation of endocytosis* (adjusted *p*-value = 0.02), *tau protein binding* (adjusted *p*-value = 1.85 × 10^-5^), and others. Among these, the regulation of *amyloid precursor protein, amyloid-beta formation*, and *tau protein binding* have been previously implicated in the pathogenesis of Alzheimer’s disease, while regulation of endocytosis is particularly important in the pathogenesis of Parkinson’s disease^22, 23^. These observations support the notion of overlapping disease-associated pathways in these common age-related neurodegenerative diseases.

## Discussion

Our analyses highlight the contributions of common and rare variants to the complex genetic architecture of LBD, a common and fatal neurodegenerative disease. Specifically, our GWAS identified five independent genome-wide significant loci (*GBA, BIN1, TMEM175, SNCA-AS1, APOE*) that influence risk for developing LBD, whereas the genome-wide gene-based aggregation tests implicated mutations in *GBA* as being critical in the pathogenesis of the disease. We further detected strong cis-eQTL colocalization signals at the *TMEM175* and *SNCA-AS1* loci, indicating that the risk of disease at these genomic regions is driven by expression changes of these particular genes. Finally, we provided definitive evidence that the risk of LBD is driven, at least in part, by the genetic variants associated with the risk of developing both Alzheimer’s disease and Parkinson’s disease.

We replicated all five GWAS signals in an independent LBD case-control dataset derived from imputed genotyping array data. Among these, *GBA* (encoding the lysosomal enzyme glucocerebrosidase), *APOE* (encoding apolipoprotein E), and *SNCA* (encoding α-synuclein) are known LBD risk genes^5–7^. In addition to these previously described loci, we identified a novel locus on chromosome 2q14.3, located 28 kb downstream to the *BIN1* gene, which is a known risk locus for Alzheimer’s disease^9^. *BIN1* encodes the bridging integrator 1 protein that is involved in endosomal trafficking. The depletion of *BIN1* reduces the lysosomal degradation of β-site APP-cleaving enzyme 1 (BACE1), resulting in increased amyloid-β production^24^. The direction of effect observed in LBD is the same as in Alzheimer’s disease (Supplementary Table 3). The observed pleiotropic effects between LBD and Alzheimer’s disease prompt us to speculate that mitigating *BIN1*-mediated endosomal dysfunction could have therapeutic implications in both neurodegenerative diseases.

A second novel LBD signal was detected within the lysosomal *TMEM175* gene on chromosome 4p16.3, a known Parkinson’s disease risk locus^10^. Deficiency of *TMEM175*, encoding a transmembrane potassium channel, impairs lysosomal function, lysosome-mediated autophagosome clearance, and mitochondrial respiratory capacity. Loss-of-function further increases the deposition of phosphorylated α-synuclein^25^, which makes *TMEM175* a plausible LBD risk gene. The direction of effect is the same in LBD as it is in Parkinson’s disease (Supplementary Table 3), and identification of *TMEM175* underscores the role of lysosomal dysfunction in the pathogenesis of Lewy body diseases.

Our data confirm the hypothesis that the LBD genetic architecture is complex and overlaps with the risk profiles of Alzheimer’s disease and Parkinson’s disease. First, several genome-wide significant risk loci in our GWAS analysis have been previously described either in the Alzheimer’s disease literature (*APOE, BIN1*) or have been associated with risk of developing Parkinson’s disease (*GBA, TMEM175, SNCA*)^9, 10, 26–28^. Second, genome-wide gene-based aggregation tests of rare mutations similarly identified *GBA*, which has been previously implicated in Parkinson’s disease^5^. Third, genetic risk scores derived from Alzheimer’s disease and Parkinson’s disease GWAS meta-analyses predicted risk for LBD independently, even after removal of the strongest signals (*APOE, GBA, SNCA,* and *LRRK2*). Interestingly, our data did not show a synergistic effect between the risk of PD and AD in the pathogenesis of LBD, though analysis of larger cohorts will be required to confirm this observation.

Comparing the patterns of the risk loci in LBD with the pattern of risk in published Parkinson’s disease and Alzheimer’s disease GWAS meta-analyses provided additional insights into this complex relationship. The directions of effect at the index variants of the *GBA* and *TMEM175* loci were the same in LBD as the directions observed in Parkinson’s disease^21^. Likewise, the directions of effect for the *BIN1* and *APOE* signals were the same as the directions detected in Alzheimer’s disease (Supplementary Table 3)^29^. However, we observed a notably different profile at the *SNCA* locus in LBD compared to PD. Our GWAS and colocalization analyses implicated *SNCA-AS1*, a non-coding RNA that regulates *SNCA* expression, as the main signal at the *SNCA* locus. In contrast, the main signal in Parkinson’s disease is detected at the 3’- end of *SNCA*^30^. This finding suggests that the regulation of *SNCA* expression may be different in LBD compared to Parkinson’s disease and that only specific *SNCA* transcripts that are regulated by *SNCA-AS1* drive risk for developing dementia. Further, *SNCA-AS1* may prove to be a more amenable therapeutic target than *SNCA* itself due to its neuronal specificity.

As part of this study, we created a foundational resource that will facilitate the study of molecular mechanisms across a broad spectrum of neurodegenerative diseases. We anticipate that these data will be widely accessed for several reasons. First, the resource is the largest whole-genome sequence repository in LBD to date. Second, the nearly 2,000 neurologically healthy, aged individuals included within this resource can be used as control subjects for the study of other neurological and age-related diseases. Third, we prioritized the inclusion of pathologically confirmed LBD patients, representing more than two thirds of the case cohort, to ensure high diagnostic accuracy among our case cohort participants. Finally, all genomes are of high quality and were generated using a uniform genome sequencing, alignment, and variant-calling pipeline. Whole genome sequencing data on this large case-control cohort has allowed us to undertake a comprehensive genomic evaluation of both common and rare variants, including immediate fine-mapping of association signals to pinpoint the functional variants at the *TMEM175* and *SNCA-AS1* loci. The availability of genome-sequence data will facilitate similar comprehensive evaluations of less commonly studied variant types, such as repeat expansions and structural variants.

Our study has limitations. We focused on individuals of European ancestry, as this is the population in which large cohorts of LBD patients were readily available. Recruiting patients and healthy controls from diverse populations will be crucial for future research to understand the genetic architecture of LBD. Another constraint is the use of short-read sequencing, rather than long-read sequencing applications, that limits the resolution of complex, repetitive, and GC-rich genomic regions^31^. Further, despite our large sample size, we had limited power to detect common genetic variants of small effect size, and additional large-scale genomic studies will be required to unravel the missing heritability of LBD.

In conclusion, our study identified novel loci as relevant in the pathogenesis of LBD. Our findings confirmed that LBD genetically intersects with Alzheimer’s disease and Parkinson’s disease and highlighted the polygenic contributions of these other neurodegenerative diseases to its pathogenesis. Determining shared molecular genetic relationships among complex neurodegenerative diseases paves the way for precision medicine and has implications for prioritizing targets for therapeutic development. We have made the whole-genome sequence data available to the research community. These genomes constitute the largest sequencing effort in LBD to date and are designed to accelerate the pace of discovery in dementia.

## Methods

### Cohort description and study design

A total of 5,154 participants of European ancestry (2,981 LBD cases, 2,173 neurologically healthy controls) were recruited across 17 European and 27 North American sites/consortia to create a genomic resource for LBD research (Supplementary Table 4). In addition to these resource genomes, we obtained convenience control genomes from (1) the Wellderly cohort (n = 1,202), a cohort of healthy, aged European-ancestry individuals recruited in the United States^32^, and (2) European-ancestry control genomes generated by the National Institute on Aging and the Accelerating Medicine Partnership - Parkinson’s Disease Initiative (www.amp-pd.org; n = 1,016). This brought the total number of control subjects available for this study to 4,391.

All control cohorts were selected based on a lack of evidence of cognitive decline in their clinical history and absence of neurological deficits on neurological examination. Pathologically confirmed control subjects (n = 605) had no evidence of significant neurodegenerative disease on histopathological examination. LBD patients were diagnosed with pathologically definite or clinically probable disease according to consensus criteria^33, 34^. The case cohort included 1,789 (69.0%) autopsy-confirmed LBD cases and 802 (31.0%) clinically probable LBD patients. 63.4% of LBD cases were male, as is typical for the LBD patient population^35^. The demographic characteristics of the cohorts are summarized in Supplementary Table 5. The appropriate institutional review boards of participating institutions approved the study (03-AG-N329, NCT02014246), and informed consent was obtained from all subjects or their surrogate decision-makers, according to the Declaration of Helsinki.

### Whole-genome sequencing

Fluorometric quantitation of the genomic DNA samples was performed using the PicoGreen dsDNA assay (Thermo Fisher). PCR-free, paired-end libraries were constructed by automated liquid handlers using the Illumina TruSeq chemistry according to the manufacturer’s protocol. DNA samples underwent sequencing on an Illumina HiSeq X Ten sequencer (v.2.5 chemistry, Illumina) using 150 bp, paired-end cycles.

### Sequence alignment, variant calling

Genome sequence data were processed using the pipeline standard developed by the Centers for Common Disease Genomics (CCDG; https://www.genome.gov/27563570/). This standard allows for whole-genome sequence data processed by different groups to generate ‘functionally equivalent’ results^36^. The GRCh38DH reference genome was used for alignment, as specified in the CCDG standard. For whole-genome sequence alignments and processing, the Broad Institute’s implementation of the functional equivalence standardized pipeline was used. This pipeline, which incorporates the GATK (2016) Best Practices^37^, was implemented in the workflow description language for deployment and execution on the Google Cloud Platform.

Single-nucleotide variants and indels were called from the processed whole-genome sequence data following the GATK Best Practices using another Broad Institute workflow for joint discovery and Variant Quality Score Recalibration. Both Broad workflows for WGS sample processing and joint discovery are publicly available (https://github.com/gatk-workflows/broad-prod-wgs-germline-snps-indels). All whole-genome sequence data were processed using the same pipeline.

### Quality control

For sample-level quality control checks, genomes were excluded from the analysis for the following reasons: (1) a high contamination rate (>5% based on VerifyBamID freemix metric)^38^, (2) an excessive heterozygosity rate (exceeding +/- 0.15 F-statistic), (3) a low call rate (≤ 95%), (4) discordance between reported sex and genotypic sex, (5) duplicate samples (determined by pi-hat statistics > 0.8), (6) non-European ancestry based on principal components analysis when compared to the HapMap 3 Genome Reference Panel (Supplementary Fig. 8)^39^, and (7) samples that were related (defined as having a pi-hat > 0.125).

For variant-level quality control, we excluded: (1) variants that showed non-random missingness between cases and controls (*p*-value ≤ 1 × 10^-4^), (2) variants with haplotype-based non-random missingness (*p*-value ≤ 1 × 10^-4^), (3) variants with an overall missingness rate of ≥ 5%, (4) non-autosomal variants (X, Y, and mitochondrial chromosomes), (5) variants that significantly departed from Hardy-Weinberg equilibrium in the control cohort (*p*-value ≤ 1 × 10^-6^), (6) variants mapping to variable, diversity, and joining (VDJ) recombination sites, as well as variants in centromeric regions +/- 10 kb (due to poor sequence alignment and incomplete resolution of the reference genome assembly at these sites)^40^, (7) variants for which the allele frequency in the aged control subjects (Wellderly cohort) significantly deviated from the other control cohorts (non-Wellderly) based on FDR-corrected chi-square tests (*p*-value < 0.05), (8) variants for which the MAFs in our control cohorts significantly differed from reported frequencies in the NHLBI Trans-Omics TOPMed database (freeze 5b; www.nhlbiwgs.org) or gnomAD (version 3.0) (FDR-corrected chi-square test *p*-value < 0.05)^41^, (9) variants that failed TOPMed variant calling filters, and (10) spanning deletions.

After these quality control filters were applied, there were 6,651 samples available for analysis. Supplementary Fig. 7 shows quality control metrics.

### Single-variant association analysis

We performed a GWAS in LBD (n = 2,591 cases and 4,027 controls) using logistic regression in PLINK (v.2.0) with a minor allele frequency threshold of >1% based on the allele frequency estimates in the LBD case cohort^42^. We used the step function in the R MASS package to determine the minimum number of principal components (generated from common single nucleotide variants) required to correct for population substructure^43^. The first two principal components in our study cohorts compared to the HapMap3 Genomic Resource Panel are shown in Supplementary Fig. 8a. Based on this analysis, we incorporated sex, age, and five principal components (PC1, PC3, PC4, PC5, PC7) as covariates in our model. Quantile-quantile plots revealed minimal residual population substructure, as estimated by the sample size-adjusted genome-wide inflation factor λ_1000_ of 1.004 (Supplementary Fig. 8b). The Bonferroni threshold for genome-wide significance was 5.0 × 10^-8^. A conditional analysis was performed for each GWAS locus by adding each respective index variant to the covariates (Supplementary Fig. 2).

For the LBD GWAS replication analysis, we obtained genotyping array data from two independent, non-overlapping, European-ancestry LBD case-control cohorts, totaling 970 LBD cases and 8,928 controls combined, as described elsewhere^44, 45^. The data were cleaned by applying the same sample- and variant-level quality control steps that were used in the discovery genomes. We imputed the data against the NHLBI TopMed imputation reference panel under default settings with Eagle v.2.4 phasing^46–48^. Variants with an R^2^ value < 0.3 were excluded. A meta-analysis of the two cohorts was performed with METAL under a fixed-effects model and variants that were significant in the discovery stage were extracted^49^.

### Colocalization analyses

Coloc (v.4.0.1) was used to evaluate the probability of LBD loci and expression quantitative trait loci (eQTLs) sharing a single causal variant^50^. This tool incorporates a Bayesian statistical framework that computes posterior probabilities for five hypotheses: namely, there is no association with either trait (hypothesis 0, H_0_); an associated LBD variant exists but no associated eQTL variant (H_1_); there is an associated eQTL variant but no associated LBD variant (H_2_); there is an association with an eQTL and LBD risk variant but they are two independent variants (H_3_); and there is a shared associated LBD variant and eQTL variant within the analyzed region (H_4_). Cis-eQTLs were derived from eQTLGen (n = 31,684 individuals; accessed February 19, 2019) and PsychENCODE (n = 1,387 individuals; accessed February 20, 2019)^16, 17^. For each locus, we examined all genes within 1Mb of a significant region of interest, as defined by our LBD GWAS (*p*-value < 5.0 × 10^-8^). Coloc was run using the default p_1_ = 10^-4^ and p_2_ = 10^-4^ priors, while the p_12_ prior was set to p_12_ = 5 × 10^-6 51^. Loci with a posterior probability for H_4_ (PPH4) ≥ 0.90 were considered colocalized. All colocalizations were subjected to sensitivity analyses to explore the robustness of our conclusions to changes in the p_12_ prior (i.e., the probability that a given variant affects both traits).

### Cell-type and tissue specificity measures

To determine specificity of a gene’s expression to a tissue or cell-type, specificity values were generated from two independent gene expression datasets: 1) bulk-tissue RNA-sequencing of 53 human tissues from the Genotype-Tissue Expression consortium (GTEx; v.8)^19^; and 2) human single-nucleus RNA-sequencing of the middle temporal gyrus from the Allen Institute for Brain Science (n = 7 cell types)^18^. Specificity values for GTEx were generated using modified code from a previous publication^52^. Expression of tissues was averaged by organ (except in the case of brain; n = 35 tissues in total). Specificity values for the Allen Institute for Brain Science-derived dataset were generated using gene-level exonic reads and the ‘generate.celltype.data’ function of the EWCE package^53^. The specificity values for both datasets and the code used to generate these values are available at https://github.com/RHReynolds/MarkerGenes.

### Heritability analysis

The narrow-sense heritability (*h^2^*), a measure of the additive genetic variance, was calculated using GREML-LDMS to determine how much of the genetic liability for LBD is explained by common genetic variants^54^. This analysis included unrelated individuals (pi-hat < 0.125, n = 2,591 LBD cases, and n = 4,027 controls) and autosomal variants with a MAF >1%. The analysis was adjusted for sex, age, and five principal components (PC1, PC3, PC4, PC5, PC7), and a disease prevalence of 0.1% to account for ascertainment bias.

### Gene-based rare variant association analysis

We conducted a genome-wide gene-based sequence kernel association test - optimized (SKAT-O) analysis of missense and loss-of-function mutations to determine the difference in the aggregate burden of rare coding variants between LBD cases and controls^11^. This analysis was performed in RVTESTS (v.2.1.0) using default parameters after annotating variants in ANNOVAR (v.2018-04/16)^55, 56^. The study cohort for this analysis consisted of 2,591 LBD cases and 4,027 control subjects. We used a MAF threshold of ≤ 1% as a filter. The covariates used in this analysis included sex, age, and five principal components (PC1, PC3, PC4, PC5, PC7). The Bonferroni threshold for genome-wide significance was 2.52 × 10^-6^ (0.05 / 19,811 autosomal genes tested).

### Predictions of LBD risk using Alzheimer’s disease and Parkinson’s disease risk scores

Genetic risk scores were generated using PLINK (v.1.9)^56^ based on summary statistics from recent Alzheimer’s disease and Parkinson’s disease GWAS meta-analyses^20, 21^. Considering the LBD cohort as our target dataset, risk allele dosages were counted across Alzheimer’s disease or Parkinson’s disease loci per sample (i.e., giving a dose of two if homozygous for the risk allele, one if heterozygous, and zero if homozygous for the alternate allele). The SNPs were weighted by their log odds ratios, giving greater weight to alleles with higher risk estimates, and a composite genetic risk score was generated across all risk loci. Genetic risk scores were z-transformed prior to analysis, centered on controls, with a mean of zero and a standard deviation of one in the control subjects. Regression models were then applied to test for association with the risk of developing LBD (based on logistic regression) or the age at death, age at onset, and disease duration (linear regression), adjusting for sex, age (risk and disease duration only), and five principal components (PC1, PC3, PC4, PC5, PC7) to account for population stratification.

### Polygenic risk score generation for pathway enrichment analysis

A genome-wide LBD polygenic risk score was generated using PRSice-2^57^. The polygenic risk score was computed by summing the risk alleles associated with LBD that had been weighted by the effect size estimates generated by performing a GWAS in the pathologically confirmed LBD cases and controls. This workflow identified the optimum *p*-value threshold (1 × 10^-4^ in our dataset) for variant selection, allowing for the inclusion of variants that failed to reach genome-wide significance but that contributed to disease risk, nonetheless. After excluding variants without an rs-identifier, the remaining 122 variants were ranked based on their GWAS *p*-values, with the *APOE*, *GBA*, *SNCA*, *BIN1* and *TMEM175* genes added to the top five positions. The list was then analyzed for pathway enrichment using the g:Profiler toolkit (v.0.1.8)^58^. We defined the genes involved in the pathways and gene sets using the following databases: (i) Gene Ontology, (ii) Kyoto Encyclopedia of Genes and Genomes, (iii) Reactome, and (iv) WikiPathways^59, 60^. Significant pathways and gene lists with a single gene or containing more than 1,000 genes were discarded. Significance was defined as a *p*-value of less than 0.05. The g:Profiler algorithm applies a Bonferroni correction to the *p*-value for each pathway to correct for multiple testing.

### Data availability

The individual-level sequence data for the resource genomes have been deposited at dbGaP (accession number: phs001963.v1.p1 NIA DementiaSeq). The GWAS summary statistics have been deposited in the GWAS catalog: https://www.ebi.ac.uk/gwas/home. eQTLGen data are available at https://www.eqtlgen.org/cis-eqtls.html. PsychENCODE QTL data are available at http://resource.psychencode.org/. Bulk-tissue RNA sequence data (GTEx version 8) are available at the Genotype-Tissue Expression consortium portal (https://www.gtexportal.org/home/).

Human single-nucleus RNA sequence data are available at the Allen Institute for Brain Science portal (portal.brain-map.org/atlases-and-data/rnaseq/human-mtg/smart-seq). Specificity values for the Allen Institute for Brain Science and GTEx data and the code used to generate these values are openly available at: https://github.com.RHReynolds/MarkerGenes.

### Code availability

Analyses were performed using open-source tools and code for analysis is available at the associated website of each software package. Genome sequence alignment and variant calling followed the implementation of the GATK Best Practices pipeline (https://github.com/gatk-workflows/broad-prod-wgs-germline-snps-indels). Contamination rates were assessed using VerifyBamID (https://genome.sph.umich.edu/wiki/VerifyBamID). Quality control checks, association analyses, and conditional analyses were performed in PLINK2 (https://www.cog-genomics.org/plink/2.0/). Data formatting and visualizations were performed in R (version 3.5.2; (https://www.r-project.org) using the following packages: MASS, tidyverse, stringr, ggrepel, data.table, viridis, ggplot2, gridExtra, grid. Imputation was performed using Eagle v.2.4 phasing (https://github.com/poruloh/Eagle). Meta-analysis was performed using METAL (https://genome.sph.umich.edu/wiki/METAL). Heritability analysis was performed using GRML-LDMS in GCTA (https://cnsgenomics.com/software/gcta). Rare variant analysis was performed using RVTESTS (v.2.1.0) (http://zhanxw.github.io/rvtests/) after annotating variant files in ANNOVAR (v.2018-04/16) (https://doc-openbio.readthedocs.io/projects/annovar/en/latest/).

Genetic risk score analyses were performed in PLINK 1.9 (https://www.cog-genomics.org/plink). LBD summary statistics were converted from hg38 to hg19 using the R implementation of the LiftOver tool, which is available from the rtracklayer package (genome.sph.umich.edu/wiki/LiftOver). Colocalization analyses were performed in R-3.2 using the packages coloc (v.4.0.1) (https://github.com/chr1swallace/coloc). Specificity values for the AIBS-derived dataset were generated using gene-level exonic reads and the ‘generate.celltype.data’ function of the EWCE package (https://github.com/NathanSkene/EWCE). Polygenic risk scores were constructed using PRSice-2 (v.2.1.1) (https://www.prsice.info). Pathway enrichment analysis was performed using the R package gprofiler2 (https://cran.r-project.org/web/packages/gprofiler2/vignettes/gprofiler2.html).

## Acknowledgments

We thank contributors who collected samples used in this study, as well as patients and families, whose help and participation made this work possible. We would like to thank Ms. Cynthia Crews for her technical assistance with DNA extractions. This research was supported in part by the Intramural Research Program of the National Institutes of Health (National Institute on Aging, National Institute of Neurological Disorders and Stroke; project numbers: 1ZIAAG000935 [PI Bryan J. Traynor, MD PhD], 1ZIANS003154 [PI Sonja W. Scholz, MD PhD], 1ZIANS0030033 and 1ZIANS003034 [David S. Goldstein, MD PhD]). Drs. Sidransky, Lopez and Tayebi were supported by the Intramural Research Program of the National Human Genome Research Institute. We would like to thank members of the International Parkinson’s Disease Genomics Consortium for providing genotyping data from 100 random Parkinson’s disease cases; these data are publicly available on dbGaP (phs00918.v1.p1). Dr. Besser gratefully acknowledges support from the National Institutes of Health (NIA K01AG063895). The American Genome Center is supported in part by an NHLBI grant: IAA-A-HL-007.001. This paper represents independent research partly funded by the National Institute for Health Research (NIHR) Biomedical Research Centre at South London and Maudsley NHS Foundation Trust and King’s College London. The study used samples from the Sant Pau Initiative on Neurodegeneration (SPIN) cohort (Sant Pau Hospital, Barcelona, Spain). We gratefully acknowledge the Knight ADRC grant P50 AG05681 (PI J.C. Morris), and its affiliated program project grant Healthy Aging and Senile Dementia P01 AG03991 (PI J.C. Morris). We would like to thank the University of Toronto Brain Bank for providing DNA specimens, and we thank the Canadian Consortium on Neurodegeneration in Aging. We acknowledge the Oxford Brain Bank, supported by the Medical Research Council, Brains for Dementia Research (Alzheimer’s Society and Alzheimer’s Research UK), Autistica UK, and the National Institute for Health Research Oxford Biomedical Research Centre. Dr. St. George-Hyslop gratefully acknowledges support from the Canadian Institutes of Research (Foundation Grant and Canadian Consortium on Neurodegeneration in Aging Grant), Wellcome Trust Collaborative Award 203249/Z/16/Z, ALS Canada Project Grant and ALS Society of Canada/Brain Canada 499553, Alzheimer Research UK, Alzheimer Society UK, and US Alzheimer Society Zenith Grant ZEN-18-529769 (PHStGH). Dr. Kevin Morgan gratefully acknowledges support by the Alzheimer’s Research UK (ARUK) Consortium. We are grateful to the Rush Alzheimer’s Disease Center for providing brain tissue and DNA samples, which was supported by the grants P30 AG10161, R01 AG15819, R01 AG17917, U01 AG46152, U01 AG61356. We thank the Dublin Brain Bank, Oregon Health & Science University Brain Bank (with support of the ADC grant 5 P30 AG008017), Duke University School of Medicine ADRC Brain Bank and Biorepository, Virginia Commonwealth University Brain Bank, Georgetown University Brain Bank, King’s College Brain Bank, Manchester University Brain Bank, NIH NeuroBioBank, the New York Brain Bank at Columbia University, and the Department of Pathology and Laboratory Medicine at the University of Kansas School of Medicine for contributing tissue specimens. We are grateful to Dr. Bernardino Ghetti from the Department of Pathology and Laboratory Medicine at Indiana University for providing tissue samples. The biospecimens from the University of California, Irvine ADRC used in this project were supported by the NIH/NIA grant P50 AG016573. Samples from the National Centralized Repository for Alzheimer’s Diseases and Related Dementias (NCRAD), which receives government support under a cooperative agreement grant (U24 AG021886) awarded by the National Institute on Aging (NIA), were used in this study. The National Alzheimer’s Coordinating Center (NACC) database is funded by the NIA/NIH Grant U01 AG016976. NACC data are contributed by the NIA-funded ADCs: P30 AG019610 (PI Eric Reiman, MD), P30 AG013846 (PI Neil Kowall, MD), P50 AG008702 (PI Scott Small, MD), P50 AG025688 (PI Allan Levey, MD, PhD), P50 AG047266 (PI Todd Golde, MD, PhD), P30 AG010133 (PI Andrew Saykin, PsyD), P50 AG005146 (PI Marilyn Albert, PhD), P50 AG005134 (PI Bradley Hyman, MD, PhD), P30 AG062677 (PI Ronald Petersen, MD, PhD), P50 AG005138 (PI Mary Sano, PhD), P30 AG008051 (PI Thomas Wisniewski, MD), P30 AG013854 (PI M. Marsel Mesulam, MD), P30 AG008017 (PI Jeffrey Kaye, MD), P30 AG010161 (PI David Bennett, MD), P50 AG047366 (PI Victor Henderson, MD, MS), P30 AG010129 (PI Charles DeCarli, MD), P50 AG016573 (PI Frank LaFerla, PhD), P50 AG005131 (PI James Brewer, MD, PhD), P50 AG023501 (PI Bruce Miller, MD), P30 AG035982 (PI Russell Swerdlow, MD), P30 AG028383 (PI Linda Van Eldik, PhD), P30 AG053760 (PI Henry Paulson, MD, PhD), P30 AG010124 (PI John Trojanowski, MD, PhD), P50 AG005133 (PI Oscar Lopez, MD), P50 AG005142 (PI Helena Chui, MD), P30 AG012300 (PI Roger Rosenberg, MD), P30 AG049638 (PI Suzanne Craft, PhD), P50 AG005136 (PI Thomas Grabowski, MD), P50 AG033514 (PI Sanjay Asthana, MD, FRCP), P50 AG005681 (PI John Morris, MD), P50 AG047270 (PI Stephen Strittmatter, MD, PhD). Data used in the preparation of this article were obtained from the AMP PD Knowledge Platform. For up-to-date information on the study, https://www.amp-pd.org. AMP PD – a public-private partnership – is managed by the FNIH and funded by Celgene, GSK, the Michael J. Fox Foundation for Parkinson’s Research, the National Institute of Neurological Disorders and Stroke, Pfizer, Sanofi, and Verily. Clinical data and biosamples used in preparation of this article were obtained from the Fox Investigation for New Discovery of Biomarkers (BioFIND), the Harvard Biomarker Study (HBS), the Parkinson’s Progression Markers Initiative (PPMI), and the Parkinson’s Disease Biomarkers Program (PDBP). BioFIND is sponsored by The Michael J. Fox Foundation for Parkinson’s Research (MJFF) with support from the National Institute of Neurological Disorders and Stroke (NINDS). The BioFIND Investigators have not participated in reviewing the data analysis or content of the manuscript. For up-to-date information on the study, visit michaeljfox.org/biofind. The Harvard NeuroDiscovery Biomarker Study (HBS) is a collaboration of HBS investigators (a full list of HBS investigators can be found at https://www.bwhparkinsoncenter.org/biobank) and funded through philanthropy and NIH and Non-NIH funding sources. The HBS Investigators have not participated in reviewing the data analysis or content of the manuscript. Data used in the preparation of this article were obtained from the Parkinson’s Progression Markers Initiative (PPMI) database (www.ppmi-info.org/data). For up-to-date information on the study, visit www.ppmi-info.org. PPMI – a public-private partnership – is funded by the Michael J. Fox Foundation for Parkinson’s Research and funding partners, including Abbvie, Allergan, Avid Radiopharmaceuticals, Biogen, BioLegend, Bristol-Myers-Squibb, Celgene, Jenali, GE Healthcare, Genentech, GlaxoSmithKline, Lilly, Lundbeck, Merck, Meso Scale Discovery, Pfizer, Piramal, Prevail Therapeutics, Roche, Sanofi Genzyme, Servier, Takeda, Teva, ucb, Verily, Voyager Therapeutics, Golub Capital. The PPMI investigators have not participated in reviewing the data analysis or content of the manuscript.

For up-to-date information on the study, visit www.ppmi-info.org. Parkinson’s Disease Biomarker Program (PDBP) consortium is supported by the National Institute of Neurological Disorders and Stroke (NINDS) and the National Institutes of Health. A full list of PDBP investigators can be found at https://pdbp.ninds.nih.gov/policy. The PDBP Investigators have not participated in reviewing the data analysis or content of the manuscript. We would like to thank the South West Dementia Brain Bank (SWDBB) for providing DNA samples for this study. The SWDBB is part of the Brains for Dementia Research Programme, jointly funded by Alzheimer’s Research UK and Alzheimer’s Society, and is supported by BRACE (Bristol Research into Alzheimer’s and Care of the Elderly) and the Medical Research Council. We are grateful to the Banner Sun Health Research Institute Brain and Body Donation Program of Sun City, Arizona, for the provision of human brain tissue (PI: Thomas G. Beach, MD). The Brain and Body Donation Program is supported by the National Institute of Neurological Disorders and Stroke (U24 NS072026 National Brain and Tissue Resource for Parkinson’s Disease and Related Disorders), the National Institute on Aging (P30 AG19610 Arizona Alzheimer’s Disease Core Center), the Arizona Department of Health Services (contract 211002, Arizona Alzheimer’s Research Center), the Arizona Biomedical Research Commission (contracts 4001, 0011, 05-901 and 1001 to the Arizona Parkinson’s Disease Consortium) and the Michael J. Fox Foundation for Parkinson’s Research. This study was supported in part by an Alzheimer’s Disease Core Center grant (P30 AG013854) from the National Institute on Aging to Northwestern University, Chicago, Illinois. We gratefully acknowledge the assistance of the Neuropathology Core. The Alzheimer’s Disease Genetics Consortium supported the collection of samples used in this study through the National Institute on Aging (NIA) grants U01 AG032984 and RC2 AG036528. This study used samples from the NINDS Repository at Coriell (https://catalog.coriell.org), as well as clinical data. Samples and data from patients included in this study were provided by the Biobank Valdecilla (PD13/0010/0024), integrated into the Spanish National Biobank Network, and they were processed following standard operating procedures with the appropriate approval of the Ethics and Scientific Committees. Data included in this manuscript has been generated with support of the grans PI16/01861 – Accion Estrategica en Salud integrated in the Spanish National I+D+P+I Plan and financed by Instituto de Salud Carlos III (ISCIII) – Subdireccion General de Evaluacion and the Fondo Europeo de Desarrollo Regional (FEDER – “Una Manera de Hacer Europa”). We thank the Mayo Clinic Brain Bank for contributing DNA samples and data from patients with Lewy body dementia, which is supported by U54 NS110435 (PI Dennis W. Dickson, MD), U01 NS100620 (PI Kejal Kantarci, MD) and The Mangurian Foundation Lewy Body Dementia Program. Sample collection and characterization is also supported by the Mayo Clinic Functional Genomics of LBD Program, NINDS R01 NS78086 (PI Owen Ross, PhD), Mayo Clinic Center for Individualized Medicine, and The Little Family Foundation. Mayo Clinic is an American Parkinson Disease Association (APDA) Mayo Clinic Information and Referral Center, an APDA Center for Advanced Research and the Mayo Clinic Lewy Body Dementia Association (LBDA) Research Center of Excellence. Dr. Zbigniew Wszolek is partially supported by the Mayo Clinic Center for Regenerative Medicine, the gifts from The Sol Goldman Charitable Trust, and the Donald G. and Jodi P. Heeringa Family, the Haworth Family Professorship in Neurodegenerative Diseases fund, and The Albertson Parkinson’s Research Foundation. We thank the Paris Neuro-CEB Brain Bank (C. Duyckaerts) for contributing DNA samples. We would like to thank the Baltimore Longitudinal Study of Aging for providing DNA samples (www.blsa.nih.gov). Tissue samples were provided by the Johns Hopkins Morris K. Udall Center of Excellence for Parkinson’s Disease Research (NIH P50 NS38377 [PI Ted M. Dawson]; U01NS097049 [PD Ted M. Dawson]) and the Johns Hopkins Alzheimer Disease Research Center (NIH P50 AG05146). This study used tissue samples and data provided by the Michigan Brain Bank, the Michigan Alzheimer’s Disease Center (P30 AG053760), and the Protein Folding Disorders Program. We are indebted to the IDIBAPS Biobank (Barcelona, Spain) for sample and data procurement. Dr. Trojanowski gratefully acknowledges support by the grant # P30 AG10124 and U19 AG062418-01A1. Tissue for this study was provided by the Newcastle Brain Tissue Resource, which is funded in part by a grant from the UK Medical Research Council (G0400074), by NIHR Biomedical Research Centre Newcastle awarded to the Newcastle upon Tyne NHS Foundation Trust and Newcastle University, and as part of the Brains for Dementia Research Programme jointly funded by Alzheimer’s Research UK and Alzheimer’s Society. Samples from the NINDS BioSend, which receives government support under a cooperative agreement grant (U24 NS095871) awarded by the National Institute of Neurological Disorders and Stroke (NINDS), were used in this study. We thank contributors who collected samples used in this study, as well as patients and their families, whose help and participation made this work possible. This study was supported in part by the Dementia with Lewy Bodies Consortium (grant #: U01 NS100610). This study acknowledges the National Institute of Neurological Disorders and Stroke (NINDS) supported Parkinson’s Disease Biomarkers Program Investigators (https://pdbp.ninds.nih.gov/sites/default/files/assets/ PDBP_investigator_list.pdf). A full list of PDBP investigators can be found at https://pdbp.ninds.nih.gov/policy. Data and biospecimens used in the preparation of this manuscript were obtained from the Parkinson’s Disease Biomarkers Program (PDBP) Consortium, part of the National Institute of Neurological Disorders and Stroke at the National Institutes of Health. Investigators include: Roger Albin, Roy Alcalay, Alberto Ascherio, Thomas Beach, Sarah Berman, Bradley Boeve, F. DuBois Bowman, Shu Chen, Alice Chen-Plotkin, William Dauer, Ted Dawson, Paula Desplats, Richard Dewey, Ray Dorsey, Jori Fleisher, Kirk Frey, Douglas Galasko, James Galvin, Dwight German, Lawrence Honig, Xuemei Huang, David Irwin, Kejal Kantarci, Anumantha Kanthasamy, Daniel Kaufer, James Leverenz, Carol Lippa, Irene Litvan, Oscar Lopez, Jian Ma, Lara Mangravite, Karen Marder, Laurie Ozelius, Vladislav Petyuk, Judith Potashkin, Liana Rosenthal, Rachel Saunders-Pullman, Clemens Scherzer, Michael Schwarzschild, Tanya Simuni, Andrew Singleton, David Standaert, Debby Tsuang, David Vaillancourt, David Walt, Andrew West, Cyrus Zabetian, Jing Zhang, and Wenquan Zou.. The PDBP Investigators have not participated in reviewing the data analysis or content of the manuscript. We thank members of the North American Brain Expression Consortium (NABEC) for providing DNA samples derived from brain tissue. Brain tissue for the NABEC cohort was obtained from the Baltimore Longitudinal Study on Aging at the Johns Hopkins School of Medicine, and from the NICHD Brain and Tissue Bank for Developmental Disorders at the University of Maryland, Baltimore, MD, USA. We would like to thank the United Kingdom Brain Expression Consortium (UKBEC) for providing DNA samples. Tissue samples and associated clinical and neuropathological data were supplied by the Parkinson’s UK Brain Bank, funded by Parkinson’s UK, a charity registered in England and Wales (258197) and in Scotland (SC037554). The authors would like to thank Parkinson’s UK for their continued support as well as the donors and family for their invaluable donation of brain tissue to the Parkinson’s UK Tissue Bank. This work was supported by a grant from the Luxembourg National Research Fund (Fonds National de Recherche, FNR) within the National Centre of Excellence in Research on Parkinson’s disease (NCER-PD). We would like to thank Estelle Sandt, Integrated Biobank Luxembourg (IBBL), and Lars Geffers, Luxembourg Centre for Systems Biomedicine (LCSB) of the University of Luxembourg, for excellent project management and the whole NCER-PD Consortium (Aguayo, Gloria; Allen, Dominic; Ammerlann, Wim; Aurich, Maike; Baldini, Federico; Balling, Rudi; Banda, Peter; Beaumont, Katy; Becker, Regina; Berg, Daniela; Betsou, Fay; Binck, Sylvia; Bisdorff, Alexandre; Bobbili, Dheeraj; Brockmann, Kathrin; Calmes, Jessica; Castillo, Lorieza; Diederich, Nico; Dondelinger, Rene; Esteves, Daniela; Ferrand, Jean-Yves; Fleming, Ronan; Gantenbein, Manon; Gasser, Thomas; Gawron, Piotr; Geffers, Lars; Giarmana, Virginie; Glaab, Enrico; Gomes, Clarissa P.C.; Goncharenko, Nikolai; Graas, Jérôme; Graziano, Mariela; Groues, Valentin; Grünewald, Anne; Gu, Wei; Hammot, Gaël; Hanff, Anne-Marie; Hansen, Linda; Hansen, Maxime; Haraldsdöttir, Hulda; Heirendt, Laurent; Herbrink, Sylvia; Hertel, Johannes; Herzinger, Sascha; Heymann, Michael; Hiller, Karsten; Hipp, Geraldine; Hu, Michele; Huiart, Laetitia; Hundt, Alexander; Jacoby, Nadine; Jarosław, Jacek; Jaroz, Yohan; Kolber, Pierre; Krüger, Rejko; Kutzera, Joachim; Landoulsi, Zied; Larue, Catherine; Lentz, Roseline; Liepelt, Inga; Liszka, Robert; Longhino, Laura; Lorentz, Victoria; Mackay, Clare; Maetzler, Walter; Marcus, Katrin; Marques, Guilherme; Martens, Jan; Mathay, Conny; Matyjaszczyk, Piotr; May, Patrick; Meisch, Francoise; Menster, Myriam; Minelli, Maura, Mittelbronn, Michel; Mollenhauer, Brit; Mommaerts, Kathleen; Moreno, Carlos; Mühlschlegel, Friedrich; Nati, Romain; Nehrbass, Ulf; Nickels, Sarah; Nicolai, Beatrice; Nicolay, Jean-Paul; Noronha, Alberto; Oertel, Wolfgang; Ostaszewski, Marek; Pachchek, Sinthuja; Pauly, Claire; Pavelka, Lukas; Perquin, Magali; Reiter, Dorothea; Rosety, Isabel; Rump, Kirsten; Sandt, Estelle; Satagopam, Venkata; Schlesser, Marc; Schmitz, Sabine; Schmitz, Susanne; Schneider, Reinhard; Schwamborn, Jens; Schweicher, Alexandra; Stallinger, Christian; Simons, Janine; Stute, Lara; Thiele, Ines; Thinnes, Cyrille; Trefois, Christophe; Trezzi, Jean-Pierre; Vaillant, Michel; Vasco, Daniel; Vyas, Maharshi; Wade-Martins, Richard; Wilmes, Paul). Finally, we would like to thank all participants of the Luxembourg Parkinson’s Study within the NCER-PD. This work was in part supported by the NIA grants U01AG049508 (PI Alison Goate, DPhil) and U01AG052411 (PI Alison Goate, DPhil). This work was supported in part by the Italian Ministry of Health (Ministero della Salute, Ricerca Sanitaria Finalizzata, grant RF-2016-02362405), the European Commission’s Health Seventh Framework Programme (FP7/2007-2013 under grant agreement 259867), the Italian Ministry of Education, University and Research (Progetti di Ricerca di Rilevante Interesse Nazionale, PRIN, grant 2017SNW5MB), the Joint Programme – Neurodegenerative Disease Research (Strength and Brain-Mend projects), granted the by Italian Ministry of Education, University and Research. This study was performed under the Department of Excellence grant of the Italian Ministry of Education, University and Research to the Rita Levi Montalcini Department of Neuroscience, University of Torino, Italy. Tissue samples were supplied by The London Neurodegenerative Diseases Brain Bank, which receives funding from the MRC (grant MR/L016397/1) and as part of the Brains for Dementia Research programme, jointly funded by Alzheimer’s Research UK and Alzheimer’s Society. DNA samples were generated by funding from the Alzheimer’s Society (422; AS-URB-18-013; PI Angela Hodges, PhD). The biospecimens from the Sunnybrook Dementia Study (https://clinicaltrials.gov/ct2/show/NCT01800214) used in this project was supported by a grant from the Canadian Institutes of Health Research (MOP13129; PIs Sandra E. Black, MD, Mario Masellis, MD, PhD) and an Early Researcher Award from the Ministry of Research, Innovation and Science (MRIS; Ontario; PI Mario Masellis, MD, PhD). The Harvard Biomarkers Study (https://www.bwhparkinsoncenter.org) is a collaborative initiative of Brigham and Women’s Hospital and Massachusetts General Hospital, co-directed by Dr. Clemens Scherzer and Dr. Bradley T. Hyman. The HBS Investigators have not participated in reviewing the current manuscript. The HBS Study Investigators are: Harvard Biomarkers Study. Co-Directors: Brigham and Women’s Hospital: Clemens R. Scherzer, Massachusetts General Hospital: Bradley T. Hyman; Investigators and Study Coordinators: Brigham and Women’s Hospital: Yuliya Kuras, Karbi Choudhury, Michael T. Hayes, Aleksandar Videnovic, Nutan Sharma, Vikram Khurana, Claudio Meleo De Gusmao, Reisa Sperling; Massachusetts General Hospital: John H. Growdon, Michael A. Schwarzschild, Albert Y. Hung, Alice W. Flaherty, Deborah Blacker, Anne-Marie Wills, Steven E. Arnold, Ann L. Hunt, Nicte I. Mejia, Anand Viswanathan, Stephen N. Gomperts, Mark W. Albers, Maria Allora-Palli, David Hsu, Alexandra Kimball, Scott McGinnis, John Becker, Randy Buckner, Thomas Byrne, Maura Copeland, Bradford Dickerson, Matthew Frosch, Theresa Gomez-Isla, Steven Greenberg, Julius Hedden, Elizabeth Hedley-Whyte, Keith Johnson, Raymond Kelleher, Aaron Koenig, Maria Marquis-Sayagues, Gad Marshall, Sergi Martinez-Ramirez, Donald McLaren, Olivia Okereke, Elena Ratti, Christopher William, Koene Van Dij, Shuko Takeda, Anat Stemmer-Rachaminov, Jessica Kloppenburg, Catherine Munro, Rachel Schmid, Sarah Wigman, Sara Wlodarcsyk; Data Coordination: Brigham and Women’s Hospital: Thomas Yi; Biobank Management Staff: Brigham and Women’s Hospital: Idil Tuncali. We thank all study participants and their families for their invaluable contributions. HBS is made possible by generous support from the Harvard NeuroDiscovery Center, with additional contributions from the Michael J Fox Foundation, NINDS U01NS082157, U01NS100603, and the Massachusetts Alzheimer’s Disease Research Center NIA P50AG005134. Data governance was provided by the METADAC data access committee, funded by ESRC, Wellcome, and MRC. (2015-2018: Grant Number MR/N01104X/1 2018-2020: Grant Number ES/S008349/1). This work made use of data and samples generated by the 1958 Birth Cohort (NCDS), which is managed by the Centre for Longitudinal Studies at the UCL Institute of Education, funded by the Economic and Social Research Council (grant number ES/M001660/1). Access to these resources was enabled via the Wellcome Trust & MRC: 58FORWARDS grant [108439/Z/15/Z] (The 1958 Birth Cohort: Fostering new Opportunities for Research via Wider Access to Reliable Data and Samples). Before 2015, biomedical resources were maintained under the Wellcome Trust and Medical Research Council 58READIE Project (grant numbers WT095219MA and G1001799). This study also used genotype and clinical data from the Wellcome Trust Case Control Consortium, and the HyperGenes Consortium. The InCHIANTI study baseline (was supported as a “targeted project” (ICS110.1/RF97.71) by the Italian Ministry of Health and in part by the United States National Institute on Aging (contracts 263 MD 9164 and 263 MD 821336), the InCHIANTI follow-up 1 (2001-2003) was funded by the United States National Institute on Aging (contracts N.1-AG-1-1 and N.1-AG-1 2111), and the InCHIANTI follow-ups 2 and 3 studies (2004-2010) were financed by the United States National Institute on Aging (contract N01-AG-5-0002). This study also used genotype and clinical data from the Wellcome Trust Case Control Consortium, and the HyperGenes Consortium. Trans-Omics in Precision Medicine (TOPMed) program imputation panel (version TOPMed-r2) is supported by the National Heart, Lung and Blood Institute (NHLB); see www.nhlbiwgs.org. TOPMed study investigators contributed data to the reference panel, which can be accessed through the Michigan Imputation Server; see https://imputationserver.sph.umich.edu. The panel was constructed and implemented by the TOPMed Informatics Research Center at the University of Michigan (3R01HL-117626-02S1; contract HHSN268201800002l). The TOPMed Data Coordinating Center (3R01HL-120393-02S1; contract HHSN268201800001l) provided additional data management, sample identify checks, and overall program coordination and support. We gratefully acknowledge the studies and participants who provided biological samples and data for TOPMed. The Molecular data for the Trans-Omics in Precision Medicine (TOPMed) program was supported by the National Heart, Lung and Blood Institute (NHLBI). Genome sequencing for “NHLBI TOPMed: Atherosclerosis Risk in Communities (ARIC)” (phs001211.v2.p2) was performed at the Broad Institute of MIT and Harvard (3R01HL092577-06S1) and at the Baylor Human Genome Sequencing Center (3U54HG003273-12S2, HHSN268201500015C). Genome sequencing for “NHLBI TOPMed: Cleveland Clinic Atrial Fibrillation (CCAF) Study” (phs001189.v1.p1) was performed at the Broad Institute of MIT and Harvard (3R01HL092577-06S1). Genome sequencing for “NHLBI TOPMed: Trans-Omics for Precision Medicine (TOPMed) Whole Genome Sequencing Project: Cardiovascular Health Study (phs001368.v1.p1) was performed at the Baylor Human Genome Sequencing Center (3U54HG003273-12S2, HHSN268201500015C). Genome sequencing for “NHLBI TOPMed: Partners HealthCare Biobank” (phs001024.v3.p1) was performed at the Broad Institute of MIT and Harvard (3R01HL092577-06S1). Genome sequencing for “NHLBI TOPMed: Whole Genome Sequencing of Venous Thromboembolism (WGS of VTE)” (phs001402.v1.p1) was performed at the Baylor Human Genome Sequencing Center (3U54HG003273-12S2, HHSN268201500015C). Genome sequencing for “NHLBI TOPMed: Novel Risk Factors for the Development of Atrial Fibrillation in Women” (phs001040.v3.p1) was performed at the Broad Institute of MIT and Harvard (3R01HL092577-06S1). Genome sequencing for “NHLBI TOPMed: The Genetics and Epidemiology of Asthma in Barbados” (phs001143.v2.p1) was performed by Illumina Genomic Services (3R01HL104608-04S1). Genome sequencing for “NHLBI TOPMed: The Vanderbilt Genetic Basis of Atrial Fibrillation” (phs001032.v4.p2) was performed at the Broad Institute of MIT and Harvard (3R01HL092577-06S1). Genome sequencing for “NHLBI TOPMed: Heart and Vascular Health Study (HVH)” (phs000993.v3.p2) was performed at the Broad Institute of MIT and Harvard (3R01HL092577-06S1) and at the Baylor Human Genome Sequencing Center (3U54HG003273-12S2, HHSN268201500015C). Genome sequencing for “NHLBI TOPMed: Genetic Epidemiology of COPD (COPDGene)” (phs000951.v3.p3) was performed at the University of Washington Northwest Genomics Center (3R01HL089856-08S1) and at the Broad Institute of MIT and Harvard (HHSN268201500014C). Genome sequencing for “NHLBI TOPMed: The Vanderbilt Atrial Fibrillation Ablation Registry” (phs000997.v3.p2) was performed at the Broad Institute of MIT and Harvard (3U54HG003067-12S2, 3U54HG003067-13S1). Genome sequencing for “NHLBI TOPMed: The Jackson Heart Study” (phs000964.v3.p1) was performed at the University of Washington Northwest Genomics Center (HHSN268201100037C). Genome sequencing for “NHLBI TOPMed: Genetics of Cardiometabolic Health in the Amish” (phs000956.v3.p1) was performed at the Broad Institute of MIT and Harvard (3R01HL121007-01S1). Genome sequencing for “NHLBI TOPMed: Massachusetts General Hospital Atrial Fibrillation (MGH AF) Study” (phs001062.v3.p2) was performed at the Broad Institute of MIT and Harvard (3R01HL092577-06S1, 3U54HG003067-12S2, 3U54HG003067-13S1, 3UM1HG008895-01S2). Genome sequencing for “NHLBI TOPMed: The Framingham Heart Study” (phs000974.v3.p2) was performed at the Broad Institute of MIT and Harvard (3U54HG003067-12S2). Core support including centralized genomic read mapping and genotype calling, along with variant quality metrics and filtering were provided by the TOPMed Informatics Research Center (3R01HL-117626-02S1; contract HHSN268201800002I). Core support including phenotype harmonization, data management, sample-identity QC, and general program coordination were provided by the TOPMed Data Coordinating Center (R01HL-120393; U01HL-120393; contract HHSN268201800001I). We gratefully acknowledge the studies and participants who provided biological samples and data for TOPMed. The Atherosclerosis Risk in Communities study has been funded in whole or in part with Federal funds from the National Heart, Lung, and Blood Institute, National Institute of Health, Department of Health and Human Services, under contract numbers (HHSN268201700001I, HHSN268201700002I, HHSN268201700003I, HHSN268201700004I, and HHSN268201700005I). The authors thank the staff and participants of the ARIC study for their important contributions. The research reported in this article was supported by grants from the National Institutes of Health (NIH) National Heart, Lung, and Blood Institute grants R01 HL090620 and R01 HL111314, the NIH National Center for Research Resources for Case Western Reserve University and Cleveland Clinic Clinical and Translational Science Award (CTSA) UL1-RR024989, the Department of Cardiovascular Medicine philanthropic research fund, Heart and Vascular Institute, Cleveland Clinic, the Fondation Leducq grant 07-CVD 03, and The Atrial Fibrillation Innovation Center, State of Ohio. This research was supported by contracts HHSN268201200036C, HHSN268200800007C, N01-HC85079, N01-HC-85080, N01-HC-85081, N01-HC-85082, N01-HC-85083, N01-HC-85084, N01-HC-85085, N01-HC-85086, N01-HC-35129, N01-HC-15103, N01-HC-55222, N01-HC-75150, N01-HC-45133, and N01-HC-85239; grant numbers U01 HL080295 and U01 HL130014 from the National Heart, Lung, and Blood Institute, and R01 AG023629 from the National Institute on Aging, with additional contribution from the National Institute of Neurological Disorders and Stroke. A full list of principal CHS investigators and institutions can be found at https://chs-nhlbi.org/pi. This manuscript was not prepared in collaboration with CHS investigators and does not necessarily reflect the opinions or views of CHS, or the NHLBI. We thank the Broad Institute for generating high-quality sequence data supported by the NHLBI grant 3R01HL092577-06S1 to Dr. Patrick Ellinor. Funded in part by grants from the National Institutes of Health, National Heart, Lung and Blood Institute (HL66216 and HL83141) and the National Human Genome Research Institute (HG04735). The Women’s Genome Health Study (WGHS) is supported by HL 043851 and HL099355 from the National Heart, Lung, and Blood Institute and CA 047988 from the National Cancer Institute, the Donald W. Reynolds Foundation with collaborative scientific support and funding for genotyping provided by Amgen. AF endpoint confirmation was supported by HL-093613 and a grant from the Harris Family Foundation and Watkin’s Foundation. The Genetics and Epidemiology of Asthma in Barbados is supported by National Institutes of Health (NIH) National Heart, Lung, Blood Institute TOPMed (R01 HL104608-S1) and: R01 AI20059, K23 HL076322, and RC2 HL101651. The research reported in this article was supported by grants from the American Heart Association to Dr. Darbar (EIA 0940116N), and grants from the National Institutes of Health (NIH) to Dr. Darbar (HL092217), and Dr. Roden (U19 HL65962, and UL1 RR024975). This project was also supported by CTSA award (UL1TR000445) from the National Center for Advancing Translational Sciences. Its contents are solely the responsibility of the authors and do not necessarily represent official views of the National Center for Advancing Translational Sciences of the NIH. The research reported in this article was supported by grants HL068986, HL085251, HL095080, and HL073410 from the National Heart, Lung, and Blood Institute. This manuscript was not prepared in collaboration with Heart and Vascular Health (HVH) Study investigators and does not necessarily reflect the opinions or views of the HVH Study or the NHLBI. This research used data generated by the COPDGene study, which was supported by NIH grants U01 HL089856 and U01 HL089897. The COPDGene project is also supported by the COPD Foundation through contributions made by an Industry Advisory Board comprised of Pfizer, AstraZeneca, Boehringer Ingelheim, Novartis, and Sunovion. Centralized read mapping and genotype calling, along with variant quality metrics and filtering were provided by the TOPMed Informatics Research Center (3R01HL-117626-02S1; contract HHSN268201800002I). Phenotype harmonization, data management, sample-identity QC, and general study coordination were provided by the TOPMed Data Coordinating Center (3R01HL-120393-02S1; contract HHSN268201800001I). We gratefully acknowledge the studies and participants who provided biological samples and data for TOPMed. This study is part of the Centers for Common Disease Genomics (CCDG) program, a large-scale genome sequencing effort to identify rare risk and protective alleles that contribute to a range of common disease phenotypes. The CCDG program is funded by the National Human Genome Research Institute (NHGRI) and the National Heart, Lung, and Blood Institute (NHLBI). Sequencing was completed at the Human Genome Sequencing Center at Baylor College of Medicine under NHGRI grant UM1 HG008898. The research reported in this article was supported by grants from the American Heart Association to Dr. Shoemaker (11CRP742009), Dr. Darbar (EIA 0940116N), and grants from the National Institutes of Health (NIH) to Dr. Darbar (R01 HL092217), and Dr. Roden (U19 HL65962, and UL1 RR024975). The project was also supported by a CTSA award (UL1 TR00045) from the National Center for Advancing Translational Sciences. Its contents are solely the responsibility of the authors and do not necessarily represent official views of the National Center for Advancing Translational Sciences or the NIH. The Jackson Heart Study (JHS) is supported and conducted in collaboration with Jackson State University (HHSN268201800013I), Tougaloo College (HHSN268201800014I), the Mississippi State Department of Health (HHSN268201800015I/HHSN26800001) and the University of Mississippi Medical Center (HHSN268201800010I, HHSN268201800011I and HHSN268201800012I) contracts from the National Heart, Lung, and Blood Institute (NHLBI) and the National Institute for Minority Health and Health Disparities (NIMHD). The authors also wish to thank the staff and participants of the JHS. The Amish studies upon which these data are based were supported by NIH grants R01 AG18728, U01 HL072515, R01 HL088119, R01 HL121007, and P30 DK072488. See publication: PMID: 18440328. The research reported in this article was supported by NIH grants K23HL071632, K23HL114724, R21DA027021, R01HL092577, R01HL092577S1, R01HL104156, K24HL105780, and U01HL65962. The research has also been supported by an Established Investigator Award from the American Heart Association (13EIA14220013) and by support from the Fondation Leducq (14CVD01). This manuscript was not prepared in collaboration with MGH AF Study investigators and does not necessarily reflect the opinions or views of the MGH AF Study investigators or the NHLBI. The Framingham Heart Study is conducted and supported by the National Heart, Lung, and Blood Institute (NHLBI) in collaboration with Boston University (Contract No. N01-HC-25195, HHSN268201500001I and 75N92019D00031). This manuscript was not prepared in collaboration with investigators of the Framingham Heart Study and does not necessarily reflect the opinions or views of the Framingham Heart Study, Boston University, or NHLBI. R.H.R was supported through the award of a Leonard Wolfson Doctoral Training Fellowship in Neurodegeneration. Raffaele Ferrari was supported by the Alzheimer’s Society (grant number 284). This work was supported by the UK Dementia Research Institute which receives its funding from DRI Ltd, funded by the UK Medical Research Council, Alzheimer’s Society and Alzheimer’s Research UK. Dr. Hardy gratefully acknowledges support from the Wellcome Trust (award number 202903/Z/16/Z), the Dolby Family Fund, National Institute for Health Research University College London Hospital Biomedical Research Centre, the BRCNIHR Biomedical Centre at University College London Hospital NHS Foundation Trust and University College London. This work was supported by the Scripps Research Translational Institute, an NIH-NCATS Clinical and Translational Science Award (CTSA; 5 UL1 RR025774). This study used the high-performance computational capabilities of the Biowulf Linux cluster at the National Institutes of Health, Bethesda, Maryland, USA (http://biowulf.nih.gov).

## AUTHOR CONTRIBUTIONS

C.L.D, B.J.T., and S.W.S. conceptualized and supervised the study. M.S.S., S.A., R.L.W., J.T.G., and Y.A. performed sample preparations; C.V. performed library preparations and genome sequencing. J.D., A.M., J.R.G., and C.L.D. performed genome sequence alignment, variant calling, and initial quality control checks. R.C., S.W.S., and B.J.T. curated the data. R.C. performed quality control checks and genome-wide association analysis, and Z.S. contributed to this analysis. R.C. also led the genome-wide gene-based rare variant analysis; M.B.M., M.D.-F., and C.B. contributed to this analysis. M.S.S. performed the heritability analysis. S.B.-C. performed the genetic risk score analysis. S.S.-A. computed polygenic risk scores and performed enrichment analyses. R.H.R., E.G. and M.R performed eQTL analyses. M.A.N. consulted on the statistical analysis. M.K.P. performed validation experiments. R.H.R., M.R., A.C., G.M., A.C., G.F. R.C.B., F.B., Z.G.-O., P.M., R.K., D.G., G.L., N.T., E.S., L.N.-K., J.-A.P., H.K, V.S., K.L.N., E.M., R.C.K., C.A.C., E.M., M.B., M.S.A., O.P., J.C.T., M.F., Q.M., E.H.B., E.R.-R., J.I., C.L., I.G.-A., P.S.-J., L.M.P., B.G., J.K., S.E.B., M.M., E.R., C.D., A.B., S.L., G.X., M.J.B., B.T., S.G., G.L., G.S., T.G.B., I.G.M., A.J.T., J.A., C.M.M., L.P., S.L., C.T., S.A.-S., A.H., D.A., G.K., S.M.K., R.W., P.P., L.M.B., J.L., L.B., A.K., A.R., A.G., D.A.B., C.S., H.R.M., R.F., S.P.-B., F.K., W.K., A.L., J.A., D.A., J.F., L.F., S.M.R., T.T., T.F., N.R.G.-R., Z.W., T.F., B.F.B., J.A.H., D.D., A.B.S., A.C., O.A.R., B.J.T., and S.W.S. provided biospecimens and clinical data. D.J.S., J.E., L.P., O.A., L.C., L.H., K.M., A.L., P.St.G-H., I.B., K.M., A.B., K.B., T.L., G.D., I.S., P.T., L.M., M.O., N.J.C., J.C.M., G.M.H., V.V.D., J.Q.T., T.G., C.S., A.C., B.J.T., and S.W.S. provided replication data. A.T., D.G.H., J.R.G., and A.B.S. provided convenience control genomes. S.W.S. wrote the initial manuscript. All authors critically reviewed and edited the article.

