## Supplementary_Materials for "Genome sequencing analysis identifies new loci associated with Lewy body dementia and provides insights into the complex genetic architecture"

Ruth Chia et al.

#### Contents


| Supplementary Information | |
|---|---|
| **Members of 'The American Genome Center'**…………………………………………... | 2 |
| **Supplementary Figures** | |
| **Supplementary Fig. 1.** Regional association plots ……………...……………………...... | 3 |
| **Supplementary Fig. 2.** Conditional analysis …………………....................................... | 5 |
| **Supplementary Fig. 3.** Sensitivity analyses …………………………………………… | 6 |
| **Supplementary Fig. 4.** GWAS variants correlate with increased *SNCA-AS1* expression | 7 |
| **Supplementary Fig. 5.** Tissue and cell-type specificity of *SNCA-AS1* and *TMEM175*... | 8 |
| **Supplementary Fig. 6.** Tissue and cell-specificity of *SNCA-AS1* and *SNCA* ………..... | 9 |
| **Supplementary Fig. 7.** Quality control metrics …………………...…..……………...... | 10 |
| **Supplementary Fig. 8.** Principal components analysis and QQ plot …....…………….. | 10 |
| **Supplementary Tables** | |
| **Supplementary Table 1.** Colocalization analysis results ……………………………... | 11 |
| **Supplementary Table 2.** Specificity values of *SNCA*, *SNCA-AS1*, *TMEM175* in GTEx and Allen Institute for Brain Science datasets ………. | 11 |
| **Supplementary Table 3.** Comparison: LBD to Alzheimer's and Parkinson's disease .. | 12 |
| **Supplementary Table 4.** Study sites and consortia ……………………………..…….. | 13 |
| **Supplementary Table 5.** Demographic characteristics …………….………....……….... | 14 |

### Supplementary Information

#### Members of 'The American Genome Center'

Anthony R. Sotis[1], Coralie Viollet[1], Gauthaman Sukumar[1], Camille Alba[1], Nathaniel Lott[1], Elisa McGrath Martinez[1], Meila Tuck[1], Jatinder Singh[1], Dagmar Bacikova[1], Xijun Zhang[1], Daniel N. Hupalo[1], Adelani Adeleye[1], Matthew D. Wilkerson[1], Harvey B. Pollard[1], Clifton L. Dalgard[1,2]

[1]The American Genome Center, Collaborative Health Initiative Research Program, Uniformed Services University of the Health Sciences, Bethesda, MD 20814, USA. [2]Department of Anatomy, Physiology & Genetics, Uniformed Services University of the Health Sciences, Bethesda, MD 20814, USA.

### Supplementary Figures

#### Supplementary Fig. 1 | Regional association plots

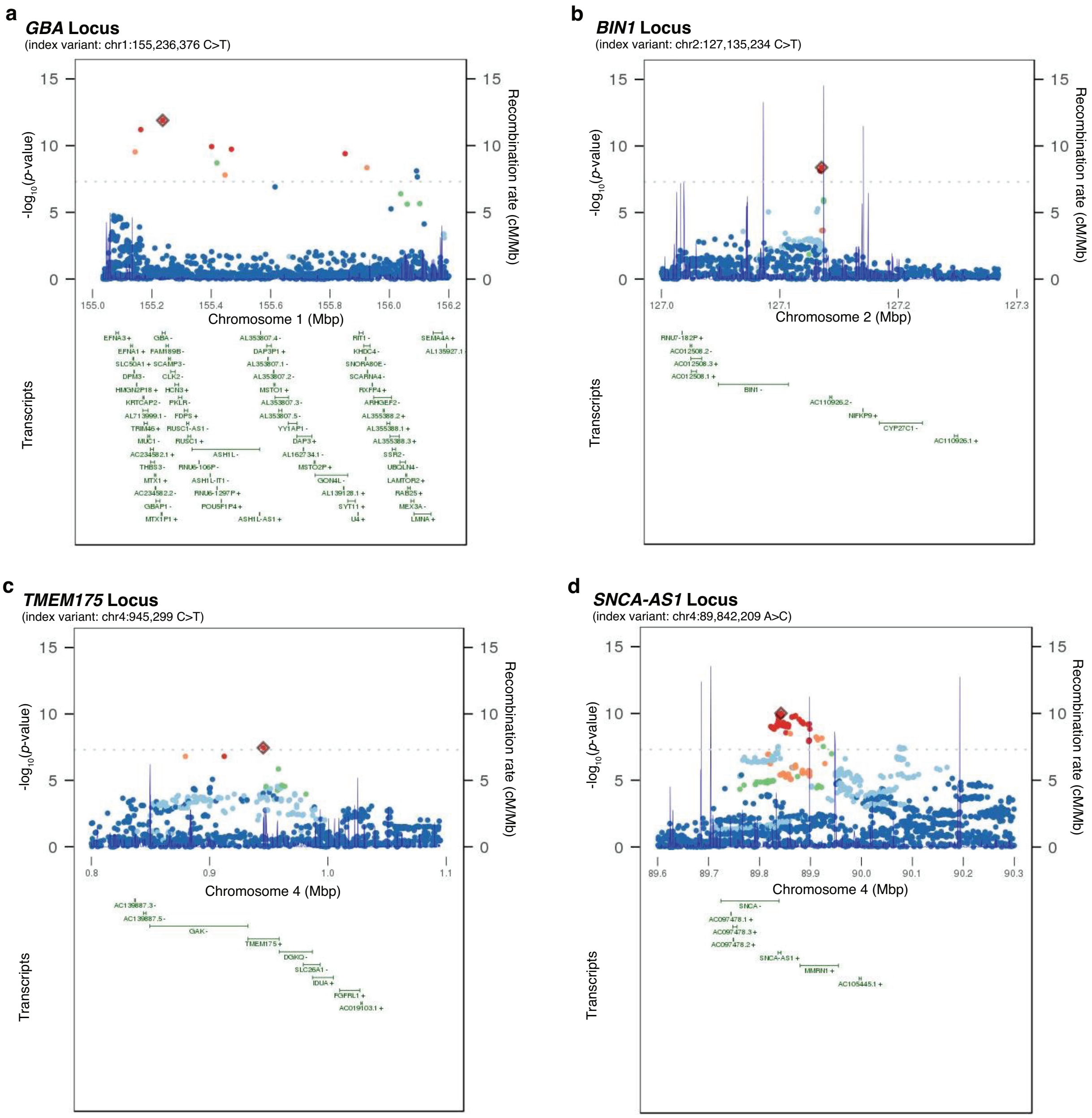


(continued on page 4)

e

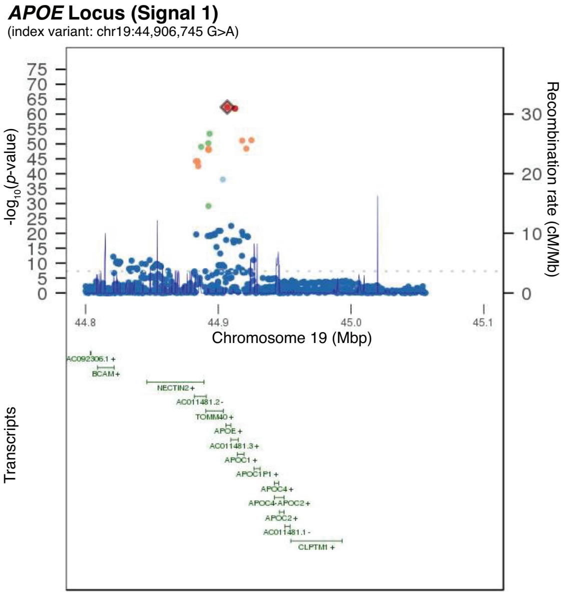


f

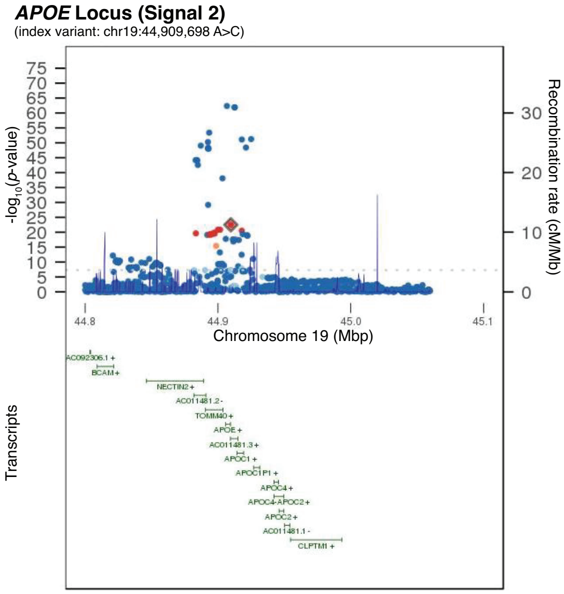


g

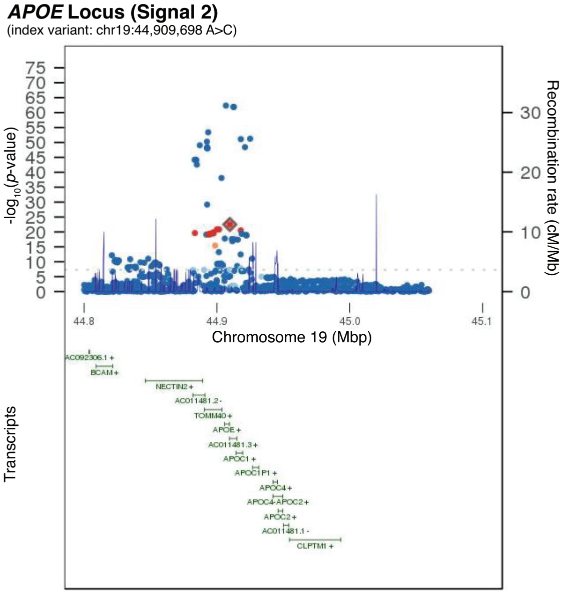


Regional association plots (**a-g**), local linkage disequilibrium, and recombination rates at the significantly associated LBD GWAS risk signals. Regional associations are plotted as a function of their genomic position, denoting the index variant by a red diamond. Single nucleotide variants or indels surrounding the index variant are color-coded to reflect the strength of linkage disequilibrium with the index variant based on pairwise $R^2$-values in the study cohort (red, 1.0 >= $R^2$ >= 0.8; orange, 0.8 > $R^2$ >= 0.6; green 0.6 > $R^2$ >= 0.4; light blue, 0.4 > $R^2$ >= 0.2; dark blue, 0.2 > $R^2$ >= 0; gray, no $R^2$ value available). Transcript annotations according to the University of California Santa Cruz genome browser are depicted under each association plot.

#### Supplementary Fig. 2 | Conditional analyses

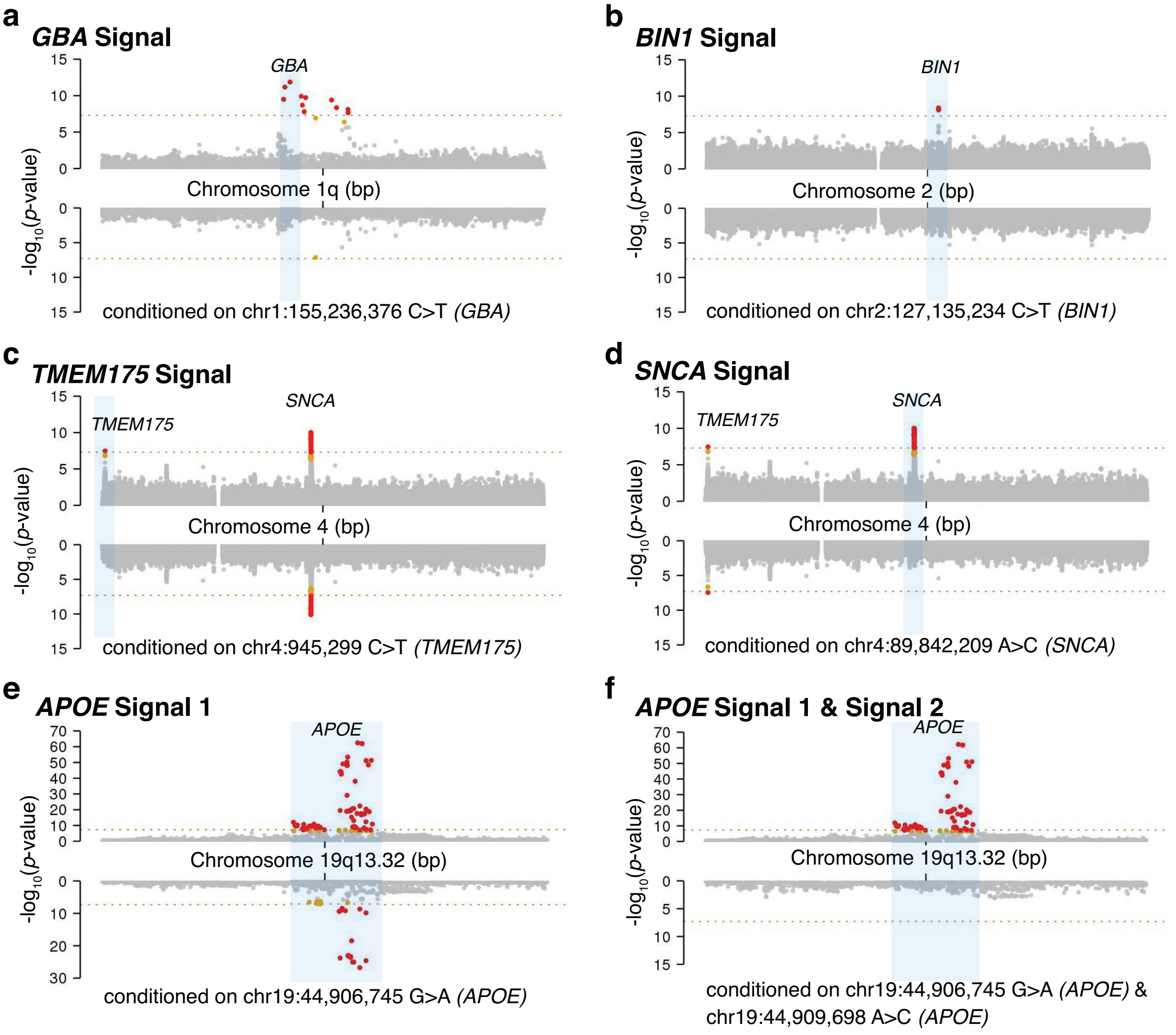


Conditional analyses for all genome-wide significant GWAS signals are depicted (**a-g**). For each panel, the x-axis denotes the chromosomal position in build 38, and the y-axis indicates the association *p*-values on a -$\log_{10}$ scale. The unconditioned GWAS signal is shown in the upper pane of each panel, while the lower pane illustrates the association results after correction for the index variant(s) at each respective signal. This analysis demonstrated two signals at the *APOE* locus [**e, f, g**]). The locus name is based on the closest gene to the index variant.

#### Supplementary Fig. 3 | Sensitivity analyses

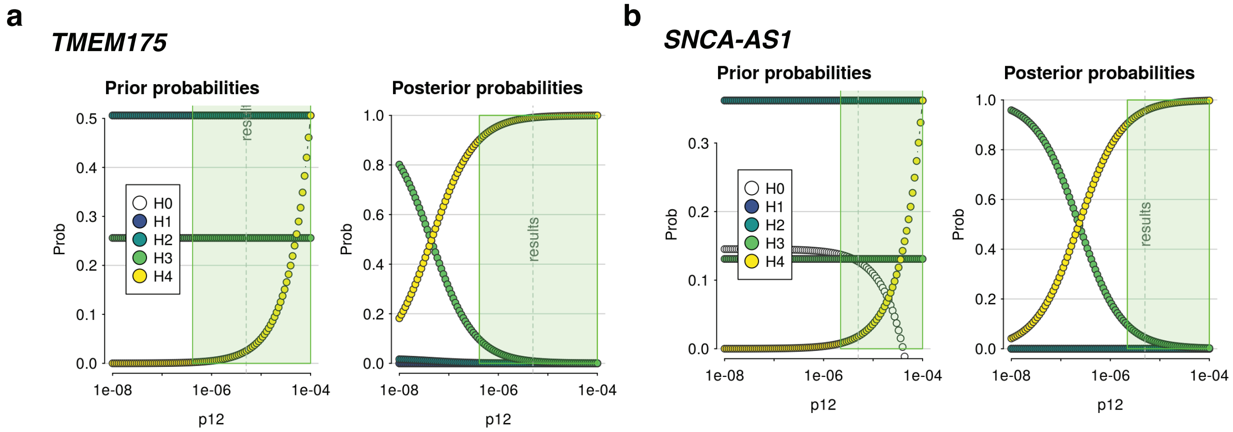


Sensitivity analyses of colocalization between eQTLs regulating **a**) *TMEM175* expression and LBD GWAS signals and **b**) *SNCA-AS1* expression and LBD GWAS signals. eQTLs for *TMEM175* were derived from eQTL-Gen, while eQTLs for *SNCA-AS1* were derived from PsychENCODE. Plots of prior (left pane) and posterior (right pane) probabilities for H0-H4 hypotheses across varying $p_{12}$ priors are shown. A dashed vertical line indicates the value of $p_{12}$ used in the initial analysis ($p_{12} = 5 \times 10^{-6}$). The green shaded areas in these plots show the regions for which the posterior probability of H4 ≥ 0.90 would still be supported. Abbreviations: H0, hypothesis 0 (no association with either trait); H1, hypothesis 1 (association with trait 1, not with trait 2); H2, hypothesis 2 (association with trait 2, not with trait 1); H3, hypothesis 3 (association with trait 1 and trait 2, two independent SNPs); H4, hypothesis 4 (association with trait 1 and trait 2, one shared SNP).

#### Supplementary Fig. 4 | GWAS variants correlate with increased *SNCA-AS1* expression

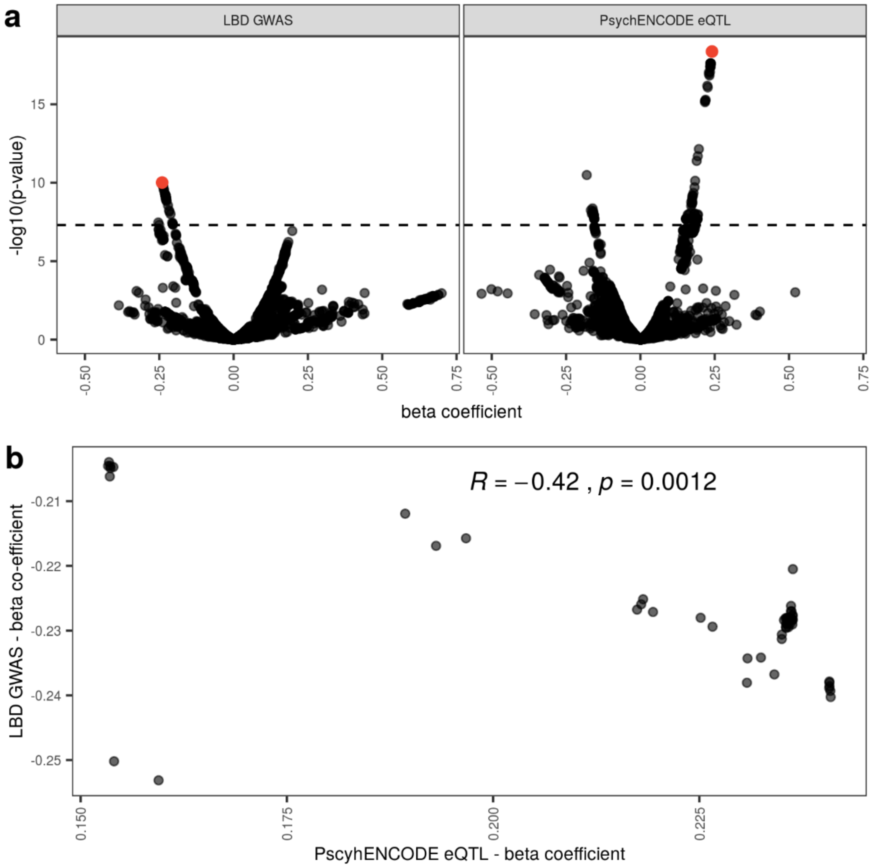


Shown here are genome-wide significant SNPs that decrease risk for LBD and their correlation with increased *SNCA-AS1* expression. **a**) Scatterplot of beta coefficients and association *p*-values (on a $-\log_{10}$ scale) for SNPs shared between the LBD GWAS (left pane) and PsychENCODE (right pane). The SNPs represented in this plot are those that are eQTLs regulating *SNCA-AS1* expression. The top SNP in the LBD GWAS (as determined by the lowest association test *p*-value) is indicated in both scatterplots by a red point. The dashed line represents the cut-off for genome-wide significance ($5 \times 10^{-8}$). **b**) Scatterplot of SNPs shared between the LBD GWAS and PsychENCODE, which pass genome-wide significance in the LBD GWAS. Spearman's rho (R) and associated *p*-value are displayed.

**Supplementary Fig. 5 | Tissue and cell-type specificity of *SNCA-AS1* and *TMEM175***

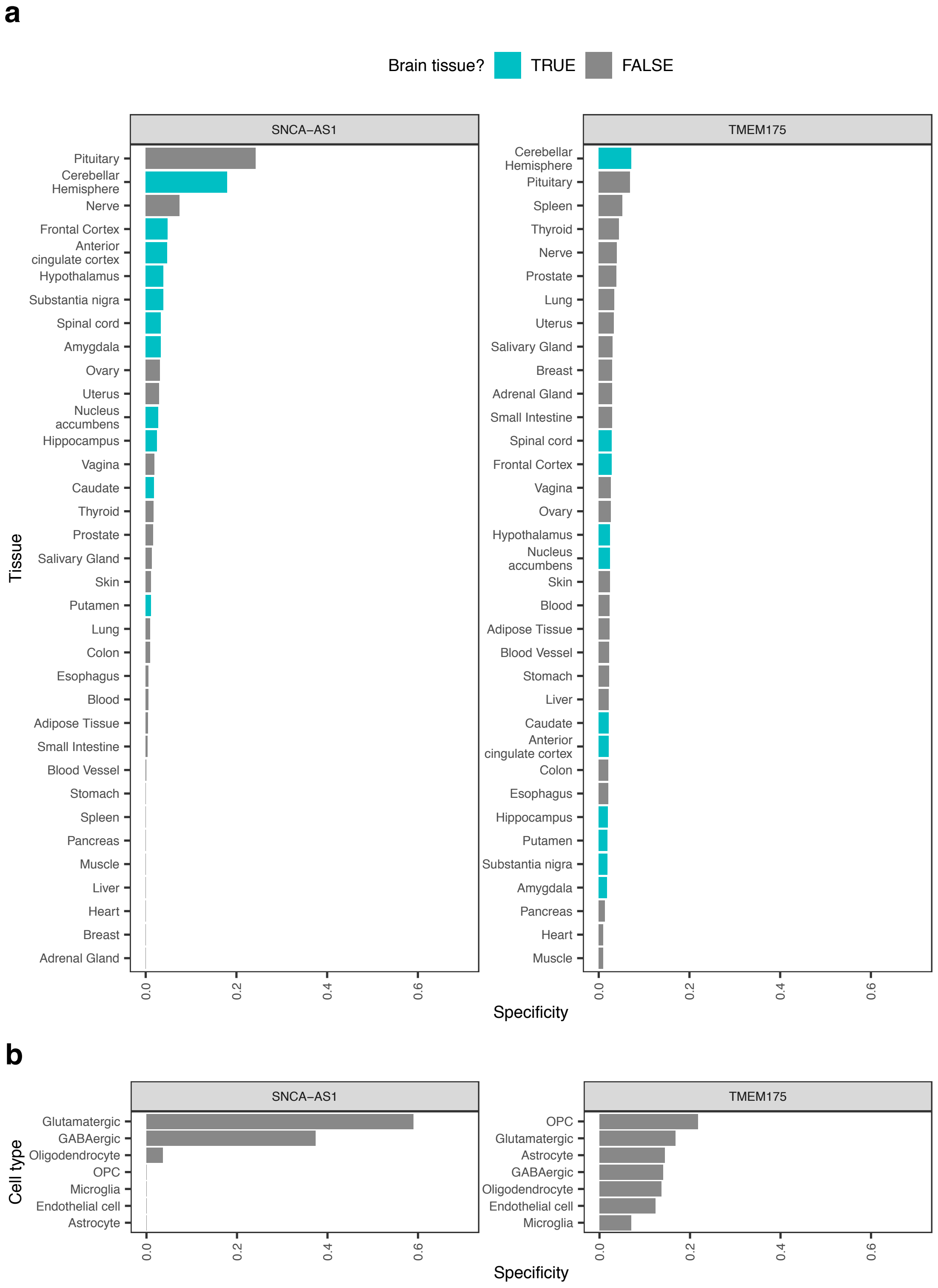


Plot of *SNCA-AS1* and *TMEM175* specificity in **a**, 35 human tissues (GTEx dataset) and **b**, seven broad categories of cell types derived from human middle temporal gyrus (Allen Institute for Brain Science dataset). Tissues are colored by whether they belong to the brain. In all plots, tissues and cell types have been ordered by specificity.

**Supplementary Fig. 6 | Tissue and cell-type specificity of *SNCA-AS1* and *SNCA***

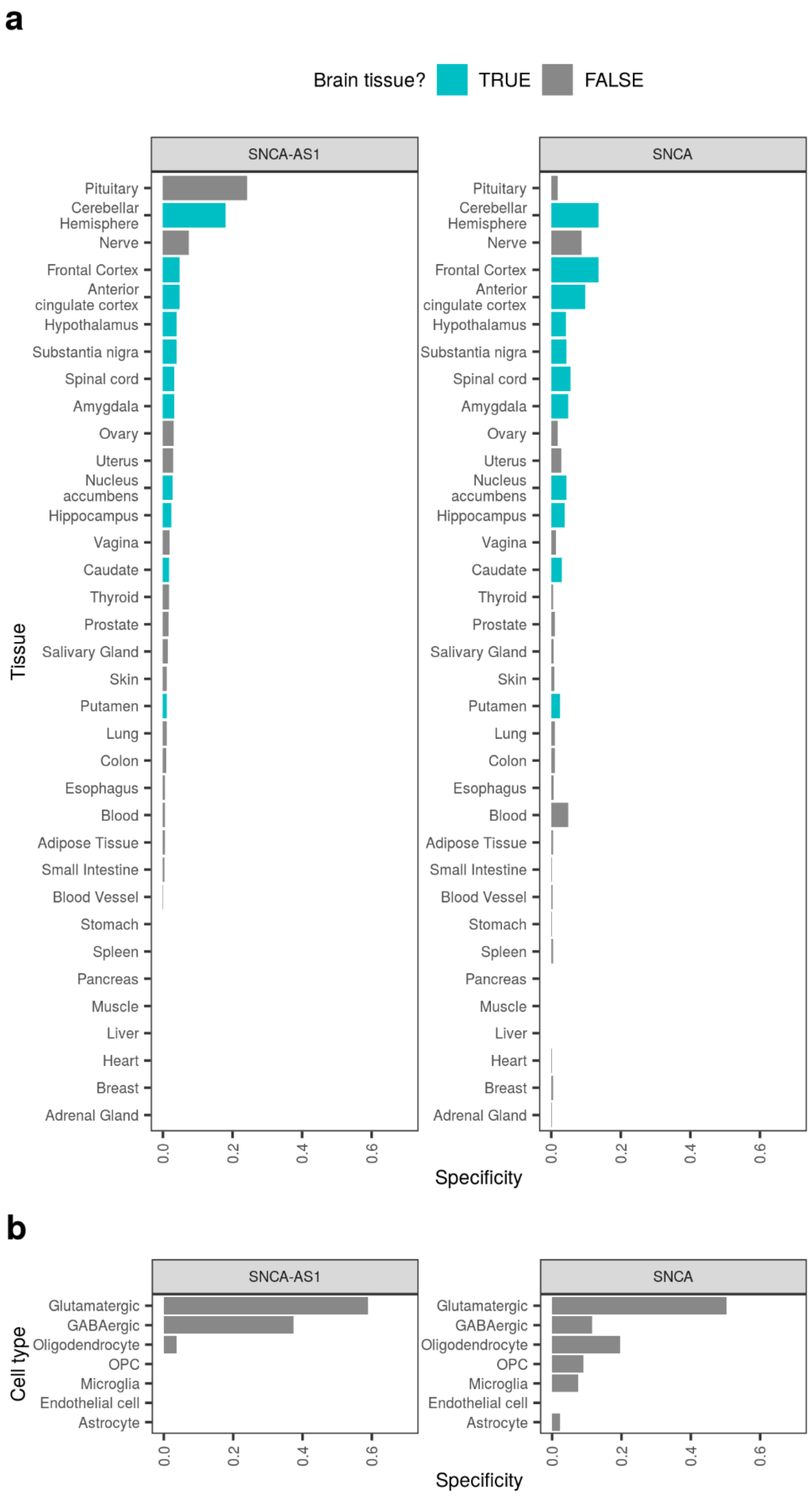


Plots of *SNCA-AS1* and *TMEM175* specificity in **a**, 35 human tissues (GTEx dataset) and **b**, seven broad categories of cell types derived from human middle temporal gyrus (Allen Institute for Brain Science dataset). Tissues are colored by whether they belong to the brain. In all plots, tissues and cell types have been ordered by specificity.

#### Supplementary Fig. 7 | Whole-genome sequence quality control metrics

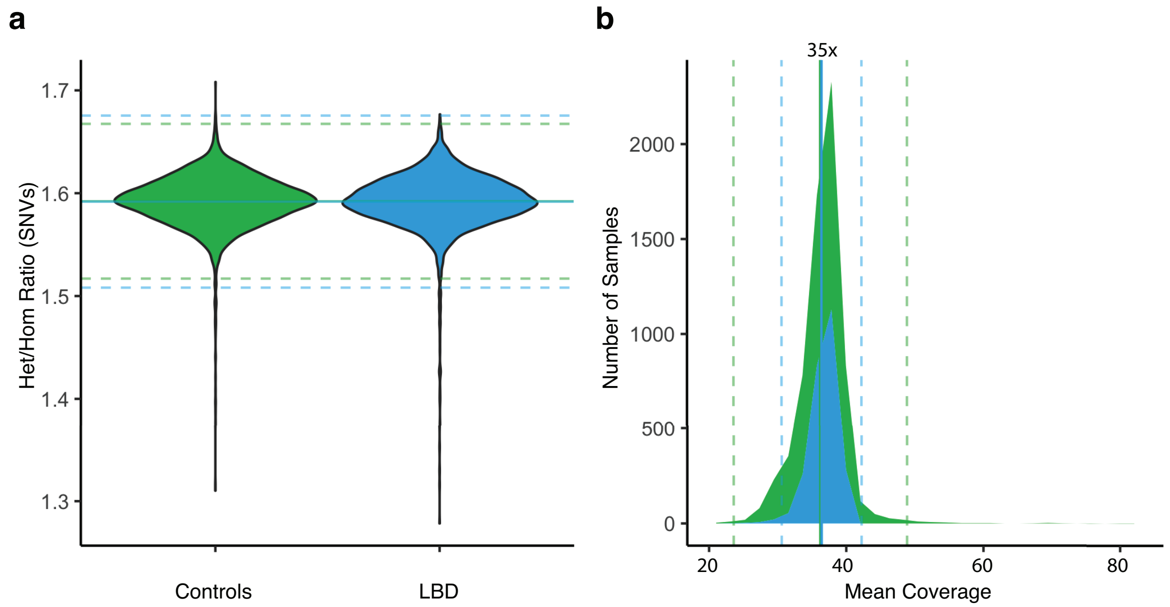


This figure depicts quality control metrics of the genome data across study cohorts. Shown are the (**a**) heterozygous-to-homozygous single nucleotide variant (SNV) ratios, and (**b**) the mean coverage across the study cohorts.

#### Supplementary Fig. 8 | Principal component analysis and QQ-plot

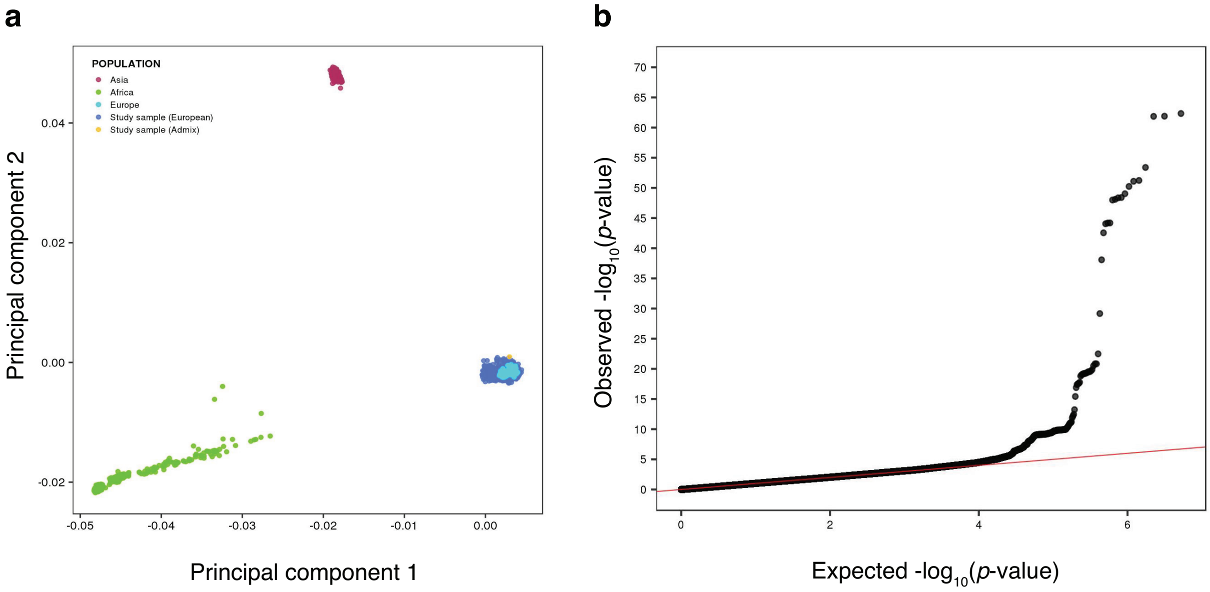


Quality control metrics of GWAS data. **a)** Population structure is shown by plotting the first two principal components of the study cohorts (n = 2,591 LBD cases and n = 4,027 controls) compared to the HapMap3 Genome Reference panel. **b)** Quantile-quantile (QQ) plot of single-variant associations depicting observed (y-axis) versus expected *p*-values (x-axis). The sample size adjusted genomic inflation factor $\lambda_{1000}$ was 1.004.

### Supplementary Tables

**Supplementary Table 1 | Colocalization analysis results**
See *SupplementaryTable1_coloc.xlsx* file

**Supplementary Table 2 | Specificity values of *SNCA*, *SNCA-AS1*, *TMEM175* in GTEx an AIBS datasets**
See *SupplementaryTable2_specificity.xlsx* file

**Supplementary Table 3.** Comparison of LBD association signals with Parkinson's and Alzheimer's disease studies

| | | | Lewy Body Dementia (Chia et al.) | | Alzheimer's Disease (Jansen et al.) | | Parkinson's Disease (Nalls et al.) | |
|---|---|---|---|---|---|---|---|---|
| **Chr.** | **Position (SNP-ID)** | **Closest Gene** | **OR (95% CI)** | ***p*-value** | **OR (95% CI)** | ***p*-value** | **OR (95% CI)** | ***p*-value** |
| 1 | 155,236,376 (rs2230288) | *GBA* | 2.89 (2.16 – 3.87) | $1.28 \times 10^{-12}$ | 1.02 (1.00 – 2.73) | $2.15 \times 10^{-2}$ | **2.19 (1.91 – 2.51)*** | **$4.11 \times 10^{-29}$** |
| 2 | 127,135,234 (rs6733839) | *BIN1* | 1.25 (1.16 – 1.35) | $4.16 \times 10^{-9}$ | **1.06 (1.05 – 2.86)** | **$1.28 \times 10^{-29}$** | 1.01 (0.97 – 1.06) | $5.75 \times 10^{-1}$ |
| 4 | 945,299 (rs6599388) | *TMEM175* | 1.25 (1.15 – 1.35) | $3.54 \times 10^{-8}$ | 1.00 (1.00 – 2.72) | $5.31 \times 10^{-2}$ | **1.20 (1.15 – 1.24)** | **$5.99 \times 10^{-21}$** |
| 4 | 89,842,209 (rs7680557) | *SNCA-AS1* | 0.79 (0.73 – 0.85) | $9.73 \times 10^{-11}$ | 1.00 (0.99 – 2.70) | $5.74 \times 10^{-2}$ | **0.95 (0.92 – 0.98)** | **$1.31 \times 10^{-3}$** |
| 19 | 44,906,745 (rs769449) | *APOE* | 2.46 (2.22 – 2.74) | $3.28 \times 10^{-72}$ | **1.19 (1.18 – 3.25)** | **0** | 0.97 (0.92 – 1.03) | $3.03 \times 10^{-1}$ |

Comparison of LBD GWAS signals with summary statistics from recent meta-analyses of genome-wide association studies in Alzheimer's disease (Jansen I et al., Nat Genet. 2019 Mar;51(3):404-413) and Parkinson's disease (Nalls M et al., Lancet Neurol. 2019 Dec;18(12):1091-1102). The SNP positions are shown according to hg38. Abbreviations: OR, odds ratio; CI, confidence interval.

**Supplementary Table 4 | Study sites/consortia that contributed samples for sequencing**

| Continent | Institution (City) | Country |
|---|---|---|
| **EUROPE** | Pitie-Salpetriere Hospital (Paris) | France |
| | University of Thessalia (Volos) | Greece |
| | Dublin Brain Bank (Dublin) | Ireland |
| | University of Torino (Torino) | Italy |
| | University of Cagliari (Cagliari) | Italy |
| | University of Bari (Bari) | Italy |
| | University of Luxembourg (Luxembourg City) | Luxembourg |
| | Hospital de Sant Pau (Barcelona) | Spain |
| | University Hospital Mutua de Terrassa (Barcelona) | Spain |
| | Biobanc-Hospital Clinic – IDIBAPS (Barcelona) | Spain |
| | Hospital Universitario "Marques de Valdecilla" (Santander) | Spain |
| | King's College London (London) | UK |
| | University College London (London) | UK |
| | Imperial College London (London) | UK |
| | University of Bristol Brain Bank (Bristol) | UK |
| | Newcastle University (Newcastle upon Tyne) | UK |
| | The University of Manchester (Manchester) | UK |
| **NORTH AMERICA** | McGill University (Montreal) | Canada |
| | University of Toronto (Toronto) | Canada |
| | Virginia Commonwealth University (Richmond, VA) | USA |
| | Banner Sun Health Research Institute (Phoenix, AZ) | USA |
| | Rush Alzheimer's Disease Center (Chicago, IL) | USA |
| | Northwestern University (Evanston, IL) | USA |
| | Parkinson's Disease Biomarker Program | USA |
| | Fox Investigation for New Discovery of Biomarkers Program | USA |
| | Indiana University School of Medicine (Indianapolis, IN) | USA |
| | National Institutes of Health (Bethesda, MD) | USA |
| | New York University Langone Medical Center (New York, NY) | USA |
| | Icahn School of Medicine at Mount Sinai (New York, NY) | USA |
| | National Cell Repository for Alzheimer's Disease (Indianapolis, IN) | USA |
| | University of California San Diego (San Diego, CA) | USA |
| | University of California (Irvine, CA) | USA |
| | North American Brain Expression Consortium | USA |
| | NINDS Biorepository at Coriell Institute (Camden, NJ) | USA |
| | University of Maryland Brain Bank (Baltimore, MD) | USA |
| | University of Kansas Medical Center (Kansas City, KS) | USA |
| | University of Michigan Brain Bank (Ann Arbor, MI) | USA |
| | Mayo Clinic (Jacksonville, FL) | USA |
| | Mayo Clinic (Rochester, MN) | USA |
| | Brigham & Women's Hospital (Boston, MA) | USA |
| | Scripps Translational Science Institute (La Jolla, CA) | USA |
| | Johns Hopkins University (Baltimore, MD) | USA |
| | Oregon Health & Science University Brain Bank (Portland, OR) | USA |
| | Baltimore Longitudinal Study on Aging (Baltimore, MD) | USA |

**Supplementary Table 5 | Demographic characteristics of study samples**

| | RESOURCE COHORTS | | CONVENIENCE CONTROLS | |
|---|---|---|---|---|
| **CHARACTERISTIC** | **LBD Cases** | **Resource Controls** | **Wellderly** | **NIA** |
| **Total number** | 2,591 | 1,908 | 1,113 | 1,006 |
| **Male, n (%)** | 1,643 (63%) | 963 (50%) | 445 (40%) | 556 (55%) |
| **Clinically ascertained, n (%)** | 802 (31%) | 1,671 (88%) | 1,113 (100%) | 638 (63%) |
| **Pathologically diagnosed, n (%)** | 1,789 (69%) | 237 (12%) | 0 (0%) | 368 (37%) |
| **Mean age, years (± s.d.)** | 75 (± 11) | 73 (± 14) | 85 (± 4) | 58 (± 19) |

All samples included in the analysis were from individuals of European ancestry. Abbreviations: s.d., standard deviation; NIA, National Institute on Aging.
